# The art of using t-SNE for single-cell transcriptomics

**DOI:** 10.1101/453449

**Authors:** Dmitry Kobak, Philipp Berens

## Abstract

Single-cell transcriptomics yields ever growing data sets containing RNA expression levels for thousands of genes from up to millions of cells. Common data analysis pipelines include a dimensionality reduction step for visualising the data in two dimensions, most frequently performed using t-distributed stochastic neighbour embedding (t-SNE). It excels at revealing local structure in high-dimensional data, but naive applications often suffer from severe shortcomings, e.g. the global structure of the data is not represented accurately. Here we describe how to circumvent such pitfalls, and develop a protocol for creating more faithful t-SNE visualisations. It includes PCA initialisation, a high learning rate, and multi-scale similarity kernels; for very large data sets, we additionally use exaggeration and downsampling-based initialisation. We use published single-cell RNA-seq data sets to demonstrate that this protocol yields superior results compared to the naive application of t-SNE.

## 1 Introduction

Recent years have seen a rapid growth of interest in single-cell RNA sequencing (scRNA-seq), or singlecell transcriptomics (Sandberg, 2014; Poulin et al., 2016). Through improved experimental techniques it has become possible to obtain gene expression data from tens of thousands of cells using full-length sequencing (Tasic et al., 2018; The Tabula Muris Consortium, 2018) and from hundreds of thousands or even millions of cells using tag-based protocols (Zeisel et al., 2018; Han et al., 2018; Saunders et al., 2018; Cao et al., 2019). Computational analysis of such data sets often entails unsupervised, exploratory steps including dimensionality reduction for visualisation. To this end, almost every study today is using t-distributed stochastic neighbour embedding, or t-SNE (van der Maaten and Hinton, 2008).

This technique maps a set of high-dimensional points to two dimensions, such that ideally, close neighbours remain close to each other and distant points remain distant. Informally, the algorithm places all points on the 2D plane, initially at random positions, and lets them interact as if they were physical particles. The interaction is governed by two ‘laws’: first, all points are repelled from each other; second, each point is attracted to its nearest neighbours (see Box 1 for a mathematical description). The most important parameter of t-SNE, called *perplexity*, controls the width of the Gaussian kernel used to compute similarities between points and effectively governs how many of its nearest neighbours each point is attracted to. The default value of perplexity in different existing implementations is 30 or 50 and the common wisdom is that “the performance of t-SNE is fairly robust to changes in the perplexity” (van der Maaten and Hinton, 2008).

### Box 1: The t-SNE algorithm

The t-SNE algorithm (van der Maaten and Hinton, 2008) is based on the SNE framework (Hinton and Roweis, 2003). SNE introduced a notion of directional similarity of point *j* to point *i*,

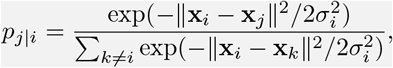

defining, for every given point *i*, a probability distribution over all points *j* ≠ *i* (all *p*_*i*|*i*_ are set to zero). The variance of the Gaussian kernel 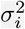 is chosen such that the perplexity of this probability distribution

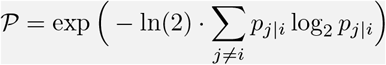

has some pre-specified value. The larger the perplexity, the larger the variance of the kernel, with the largest possible perplexity value equal to *n* − 1 corresponding to 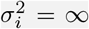 and the uniform probability distribution (*n* is the number of points in the data set). Importantly, for any given perplexity value 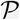, all but 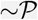 nearest neighbours of point *i* will have *p*_*j*|*i*_ very close to zero. For mathematical and computational convenience, *symmetric SNE* defined undirectional similarities

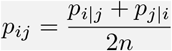

such that 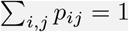, i.e. this is a valid probability distribution on the set of all pairs (*i, j*).

The main idea of SNE and its modifications is to arrange the *n* points in a low-dimensional space such that the similarities *q_ij_* between low-dimensional points match *p_ij_* as close as possible in terms of the Kullback-Leibler divergence. The loss function is thus

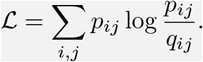

The main idea of t-SNE was to use a t-distribution with one degree of freedom (also known as Cauchy distribution) as the low-dimensional similarity kernel:

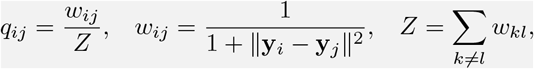

where **y**_*i*_ are low-dimensional coordinates (and *q_ii_* = 0). As a matter of definition, we consider any method that uses the t-distribution as the output kernel and Kullback-Leibler divergence as the loss function to be “t-SNE”; similarities *p*_*j*|*i*_ can in principle be computed using non-Euclidean distances instead of ║**x**_*i*_ − **x**_*j*_║ or can use non-perplexity-based calibrations.

To justify our intuitive explanation in terms of attractive and repulsive forces, we can rewrite the loss function as follows:

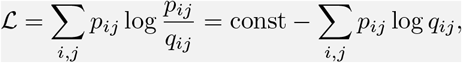

and dropping the constant,

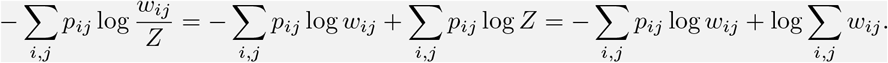

To minimise 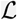, the first sum should be as large possible, which means large *w_ij_*, i.e. small ║**y**_*i*_ − **y**_*j*_║, meaning an attractive force between points *i* and *j* whenever *p_ij_* ≠ 0. At the same time, the second term should be as small as possible, meaning small *w_ij_* and a repulsive force between any two points *i* and *j*, independent of the value of *p_ij_*.

When applied to high-dimensional but well-clustered data, t-SNE tends to produce a two-dimensional visualisation with distinctly isolated clusters, which in many cases are in good agreement with the clusters produced by a dedicated clustering algorithm. This attractive property as well as the lack of serious competitors until very recently (McInnes et al., 2018; Becht et al., 2019) made t-SNE the de facto standard for visual exploration of scRNA-seq data. At the same time, t-SNE has well known, but frequently overlooked weaknesses (Wattenberg et al., 2016). Most importantly, it often fails to preserve the global geometry of the data. This means that when t-SNE places cells into several distinct clusters, the relative position of these clusters on the 2D plot is almost arbitrary and depends on random initialisation more than on anything else. While this may not be a problem in some situations, scRNA-seq data sets often exhibit biologically meaningful hierarchical structure, e.g. encompass several very different cell classes, each further divided into various types. Typical t-SNE plots do not capture such global structure, yielding a suboptimal and potentially misleading visualisation. In our experience, the larger the data set, the more severe this problem becomes. Other notable challenges include performing t-SNE visualisations for very large data sets (e.g. a million of cells or more), or mapping cells collected in follow-up experiments onto an existing t-SNE visualisation.

Here we explain how to achieve improved t-SNE visualisations that preserve the global geometry of the data. Our method relies on several modifications of the standard pipeline, such as providing PCA initialisation, employing multi-scale similarities (Lee et al., 2015; De Bodt et al., 2018), increasing the learning rate (Belkina et al., 2018), and for very large data sets, additionally using so called exaggeration and downsampling-based initialisation. To demonstrate these techniques we use a published Smart-seq2 data set with 24 thousand cells and several UMI-based data sets with up to two million cells (Table 1). We used FIt-SNE (Linderman et al., 2019), a recently developed fast t-SNE implementation, for all experiments.

**Table 1:**
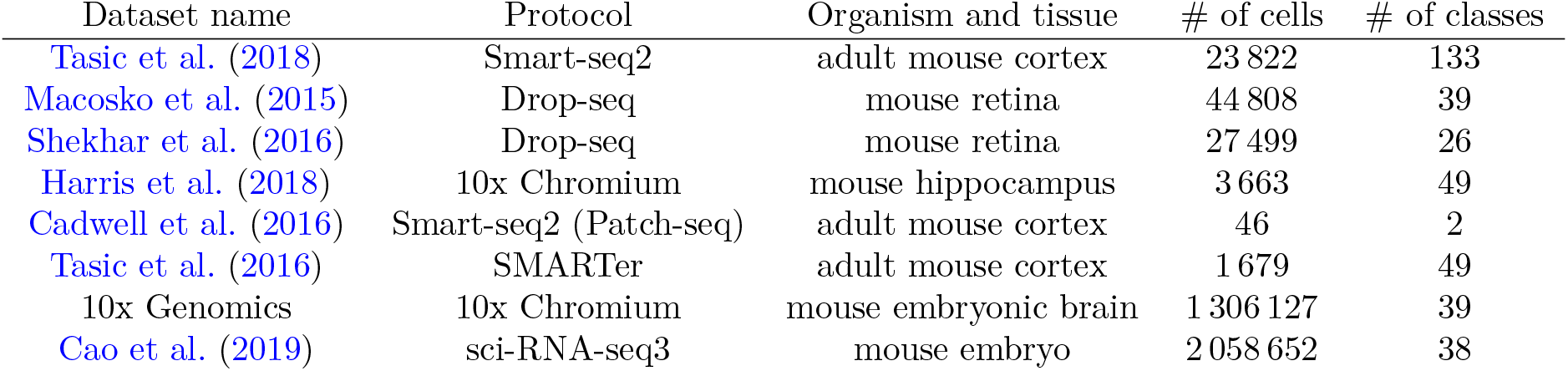
Data sets used in this study, listed in the order of appearance in the text. In all cases, we relied on quality control and clustering performed in the original publications. Cell numbers are after quality control. For the 10x Genomics data set we used the cluster labels from Wolf et al. (2018).

| Dataset name | Protocol | Organism and tissue | # of cells | # of classes |
| --- | --- | --- | --- | --- |
| Tasic et al. (2018) | Smart-seq2 | adult mouse cortex | 23 822 | 133 |
| Macosko et al. (2015) | Drop-seq | mouse retina | 44 808 | 39 |
| Shekhar et al. (2016) | Drop-seq | mouse retina | 27 499 | 26 |
| Harris et al. (2018) | 10x Chromium | mouse hippocampus | 3 663 | 49 |
| Cadwell et al. (2016) | Smart-seq2 (Patch-seq) | adult mouse cortex | 46 | 2 |
| Tasic et al. (2016) | SMARTer | adult mouse cortex | 1 679 | 49 |
| 10x Genomics | 10x Chromium | mouse embryonic brain | 1 306 127 | 39 |
| Cao et al. (2019) | sci-RNA-seq3 | mouse embryo | 2 058 652 | 38 |

In many challenging cases, our t-SNE pipeline yields visualisations that are better than the state of the art. We discuss its advantages and disadvantages compared to UMAP (McInnes et al., 2018), a recent dimensionality reduction method that is gaining popularity in the scRNA-seq community (Becht et al., 2019). We also describe how to position new cells on an existing t-SNE reference atlas and how to visualise multiple related data sets side by side in a consistent fashion. We focused on singlecell transcriptomics but our recommendations are more generally applicable to any data set that has hierarchical organisation, which is often the case e.g. in single-cell flow or mass cytometry (Amir et al., 2013; Unen et al., 2017; Belkina et al., 2018) or whole-genome sequencing (Li et al., 2017; Diaz-Papkovich et al., 2018), as well as outside of the biology (Schmidt, 2008).

## 2 Results

### 2.1 Preserving global geometry with t-SNE: synthetic data

To illustrate that the default t-SNE tends to misrepresent the global geometry, we first consider a toy example (Figure 1). This synthetic data set consists of points sampled from fifteen 50-dimensional spherical Gaussian distributions (‘cell types’), grouped into three broad ‘classes’. The data are generated such that the three classes are very distinct and non-overlapping; the types within two classes (*n* = 100 and *n* = 1000 per type respectively) do not overlap either, and the types within the third class (*n* = 2000 per type) are partially overlapping. As a result, this data set exhibits hierarchical structure, typical for scRNA-seq data.

**Figure 1:**
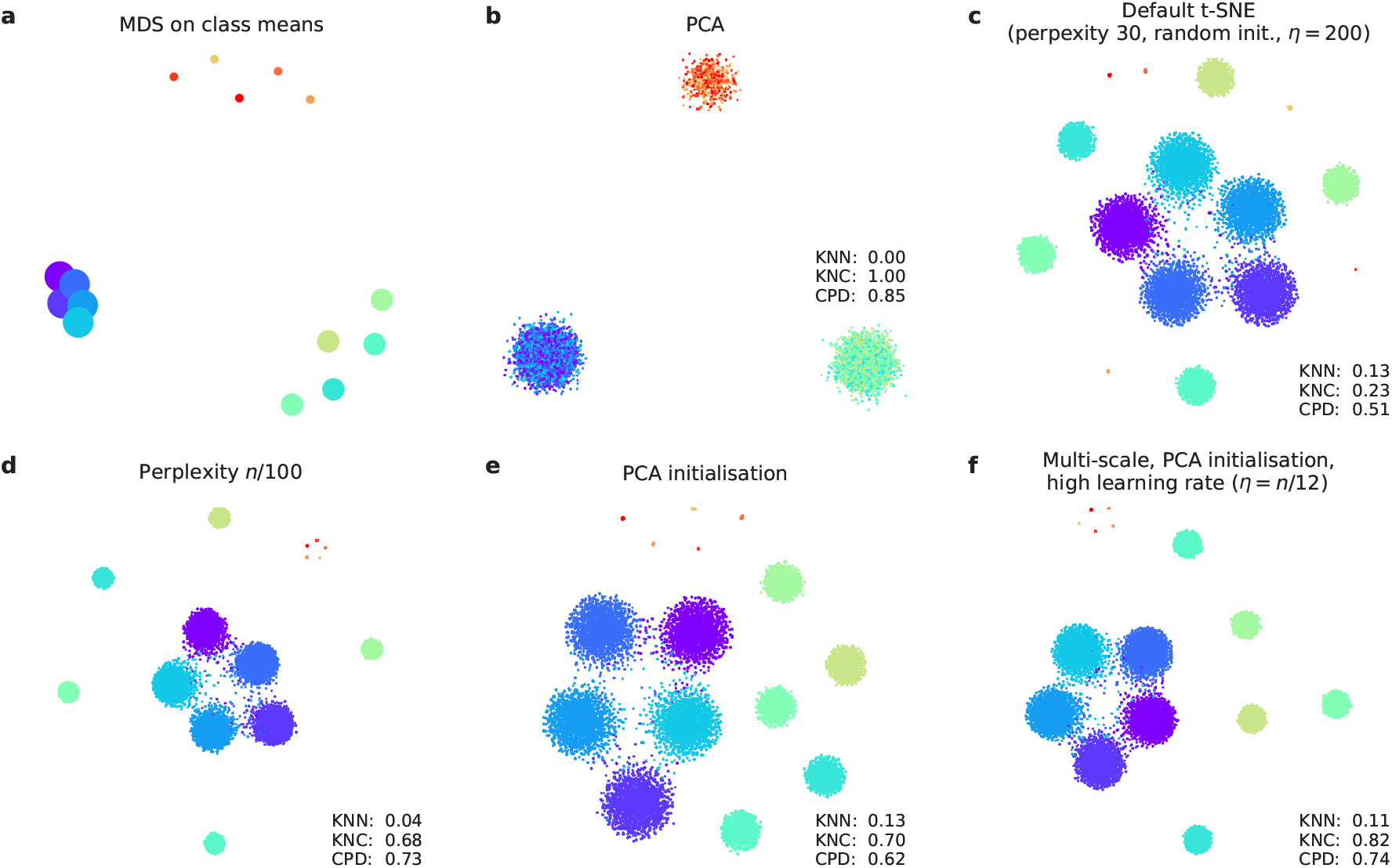
Synthetic data set. The points were sampled from a mixture of fifteen 50-dimensional Gaussian distributions. Total sample size *n* = 15 500. **(a)** Multidimensional scaling of 15 class means. Point sizes are proportional to the number of points per class. **(b)** The first two principal components of the data. Point colours denote class membership. KNN: 10-nearest neighbour preservation, KNC: 4-nearest classes preservation, CPD: Spearman correlation between pairwise distances. **(c)** Default t-SNE with perplexity 30, random initialisation, and learning rate 200. **(d)** T-SNE with perplexity *n*/100 = 155. **(e)** T-SNE with PCA initialisation. **(f)** T-SNE with multi-scale similarities (perplexity combination of 30 and *n*/100 = 155), PCA initialisation, and learning rate *n*/12 ≈ 1300.

Two classical methods to visualise high-dimensional data are multidimensional scaling (MDS) and principal component analysis (PCA). MDS is difficult to compute with a large number of points (here *n* = 15 500), but can be easily applied to class means (*n* = 15), clearly showing the three distinct classes (Figure 1a). PCA can be applied to the whole data set and demonstrates the same large-scale structure of the data (Figure 1b), but no within-class structure can be seen in the first two PCs. In contrast, t-SNE clearly shows all 15 types, correctly displaying ten of them as fully isolated and five as partially overlapping (Figure 1c). However, the isolated types end up arbitrarily placed, with their positions mostly depending on the random seed used for initialisation.

Our recipe for a more faithful t-SNE visualisation is based on three modifications that have been previously suggested in various contexts: multi-scale similarities (Lee et al., 2015; De Bodt et al., 2018), PCA initialisation, and increased learning rate (Belkina et al., 2018). In order to quantify numerically the quality, or ‘faithfulness’, of a given embedding, we used three different metrics:

**KNN** The fraction of *k*-nearest neighbours in the original high-dimensional data that are preserved as *k*-nearest neighbours in the embedding (Lee and Verleysen, 2009). We used *k* = 10 and computed the average across all *n* points. This measure quantifies the local, or ‘microscopic’ structure.
**KNC** The fraction of *k*-nearest class means in the original data that are preserved as *k*-nearest class means in the embedding. This is computed for class means only and averaged across all classes. For the synthetic data set we used *k* = 4, and for the real data sets analysed below we used *k* = 10. This measure quantifies the ‘mesoscopic’ structure.
**CPD** Spearman correlation between pairwise distances in the high-dimensional space and in the embedding (Becht et al., 2019). Computed across all pairs among 1000 randomly chosen points. Quantifies the global, or ‘macroscropic’ structure.

Applying these metrics to the PCA and t-SNE embeddings (Figure 1b,c) shows that t-SNE is much better than PCA in preserving the local structure (KNN 0.13 vs. 0.00) but much worse in preserving the global structure (KNC 0.23 vs. 1.00 and CPD 0.51 vs. 0.85).

Figure 1c used perplexity 30, which is the default value in most t-SNE implementations. Much larger values can yield qualitatively different outcomes. As large perplexity yields longer-ranging attractive forces during t-SNE optimisation, the visualisation loses some fine detail but pulls larger structures together. As a simple rule of thumb, we take 1% of the sample size as a ‘large’ perplexity for any given data set; this corresponds to perplexity 155 for our simulated data and results in five small clusters belonging to the same class attracted together (Figure 1d). Our metrics confirmed that, compared to the standard perplexity value, the local structure (KNN) deteriorates but the global structure (KNC and CPD) improves. To obtain multi-scale similarities and preserve both local and global structure, it has been suggested to use multiple perplexity values at the same time (Lee et al., 2015; De Bodt et al., 2018). We adopt this approach in our final pipeline and, whenever *n*/100 ≫ 30, combine perplexity 30 with the ‘large’ perplexity *n*/100 (see below; separate evaluation not shown here).

Another approach to preserve global structure is to use an informative initialisation, e.g. the first two PCs (after appropriate scaling, see Methods). This ‘injects’ the global structure into the t-SNE embedding which is then preserved during the course of t-SNE optimisation while the algorithm optimises the fine structure (Figure 1e). Indeed, KNN did not depend on initialisation, but both KNC and CPD markedly improved when using PCA initialisation. PCA initialisation is also convenient because it makes the t-SNE outcome reproducible and not dependent on a random seed.

The third ingredient in our t-SNE protocol is to increase the learning rate. The default learning rate in most t-SNE implementations is *η* = 200 which is not enough for large data sets and can lead to poor convergence and/or convergence to a suboptimal local minimum (Belkina et al., 2018). A recent Python library for scRNA-seq analysis, `scanpy`, increased the default learning rate to 1000 (Wolf et al., 2018), whereas Belkina et al. (2018) suggested to use *η* = *n*/12. We adopt the latter suggestion and use *η* = *n*/12 whenever it is above 200. This does not have a major influence on our synthetic data set (because its sample size is not large enough for this to matter), but will be important later on.

Putting all three modifications together, we obtain the visualisation shown in Figure 1f. The quantitative evaluation confirmed that in terms of the mesoscopic/macroscopic structure, our suggested pipeline strongly outperformed the default t-SNE and was better than large perplexity or PCA initialisation on their own. At the same time, in terms of the miscroscopic structure, it achieved a compromise between the small and the large perplexities.

### 2.2 Preserving global geometry with t-SNE: real data

To demonstrate these ideas on a real-life data set, we chose to focus on the data from Tasic et al. (2018). It encompasses 23 822 cells from adult mouse cortex, split by the authors into 133 clusters with strong hierarchical organisation. Here and below we used a standard preprocessing pipeline consisting of library size normalisation, feature selection, log-transformation, and reducing the dimensionality to 50 PCs (see Methods).

In the Tasic et al. data, three well-separated groups of clusters are apparent in the MDS (Figure 2a) and PCA (Figure 2b) plots, corresponding to excitatory neurons (cold colours), inhibitory neurons (warm colours), and non-neural cells such as astrocytes or microglia (grey/brown colours). Performing PCA on these three data subsets separately (Figure S1) reveals further structure inside each of them: e.g. inhibitory neurons are well separated into two groups, *Pvalb/SSt*-expressing (red/yellow) and *Vip/Lamp5*-expressing (purple/salmon), as can also be seen in Figure 2a. This demonstrates the hierarchical organisation of the data.

**Figure 2:**
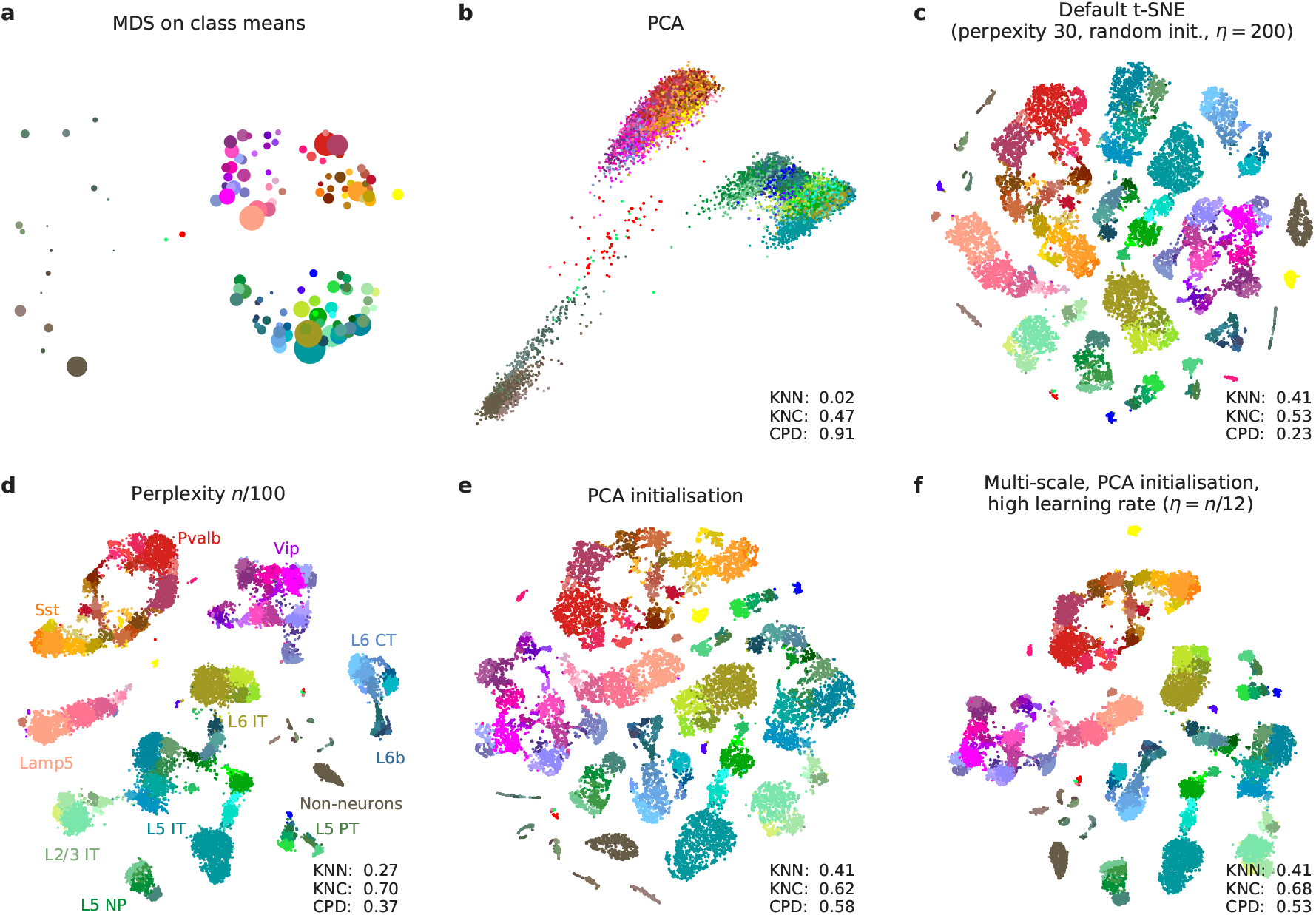
Tasic et al. data set. Sample size *n* = 23 822. Cluster assignments and cluster colours are taken from the original publication (Tasic et al., 2018). Warm colours correspond to inhibitory neurons, cold colours correspond to excitatory neurons, brown/grey colours correspond to non-neural cells. **(a)** MDS on class means (*n* = 133). Point sizes are proportional to the number of points per class. **(b)** The first two principal components of the data. KNN: 10-nearest neighbour preservation, KNC: 10-nearest classes preservation, CPD: Spearman correlation between pairwise distances. **(c)** Default t-SNE with perplexity 30, random initialisation, and learning rate 200. **(d)** T-SNE with perplexity *n*/100 = 238. Labels denote large groups of clusters. **(e)** T-SNE with PCA initialisation. **(f)** T-SNE with multi-scale similarities (perplexity combination of 30 and *n*/100 = 238, PCA initialisation, and learning rate *n*/12 ≈ 2000.

This global structure is missing from a standard t-SNE visualisation (Figure 2c): excitatory neurons, inhibitory neurons, and non-neural cells are all split into multiple ‘islands’ that are shuffled among each other. For example, the group of purple clusters (*Vip* interneurons) is separated from a group of salmon clusters (a closely related group of *Lamp5* interneurons) by some excitatory clusters, misrepresenting the hierarchy of cell types. This outcome is not a strawman: Tasic et al. (2018) features a t-SNE figure qualitatively very similar to our visualisation. Perplexity values in the common range (e.g. 20, 50, 80) yield similar results, confirming that t-SNE is not very sensitive to the exact value of perplexity.

In contrast, setting perplexity to 1% of the sample size, in this case to 238, pulls together large groups of related types, improving the global structure (KNC and CPD increase), at the expense of losing some of the fine structure (KNN decreases, Figure 2d). PCA initialisation with default perplexity also improves the global structure (KNC and CPD increase, compared to the default t-SNE, Figure 2e). Finally, our suggested pipeline with multi-scale similarities (perplexity combination of 30 and *n*/100 = 238), PCA initialisation, and learning rate *n*/12 ≈ 2000 yields an embedding with high values of all three metrics (Figure 2f). Compared to the default parameters, these settings slowed down FIt-SNE from ~30 s to ~2 m, which is nevertheless still an acceptable time.

It is instructive to study systematically how the choice of parameters influences the embedding quality (Figure 3). We found that the learning rate only influences KNN: the higher the learning rate, the better preserved is the local structure, until is saturates at around *n*/12 (Figure 3a), in agreement with the results of Belkina et al. (2018). The other two metrics, KNC and PDC, are not affected by the learning rate (Figure 3c,e). The perplexity controls the trade-off between KNN and KNC: the higher the perplexity combined with 30, the worse is the microscropic structure (Figure 3b) but the better is the mesoscopic structure (Figure 3d). Our choice of *n*/100 provides a reasonable compromise. Finally, the PCA initialisation strongly improves the macroscopic structure as measured by the PDC (Figure 3e,f), while the other two parameters have little influence on it.

**Figure 3:**
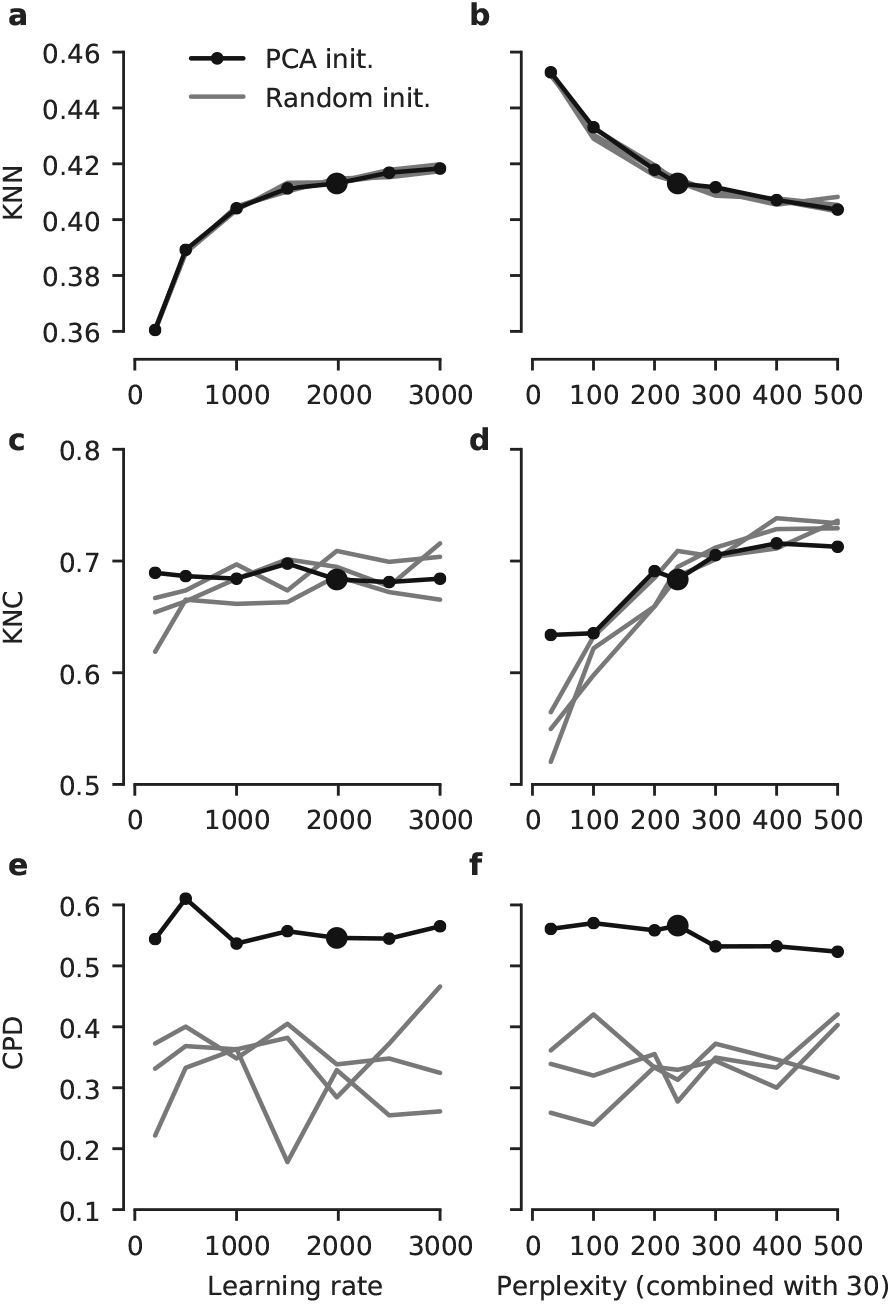
The influence of parameter values on embedding quality. All panels show quality assessments of various t-SNE embeddings of the Tasic et al. data set. **(a)** 10-nearest neighbours preservation (KNN) as a function of learning rate. Black line shows PCA initialisation, grey lines show random initialisations with three different random seeds. Large black dot denotes our preferred parameter values. Perplexity combination of 30 and *n*/100. **(b)** 10-nearest neighbours preservation as a function of perplexity used in combination with perplexity 30. Learning rate *n*/12. **(c–d)** The same for 10-nearest classes preservation (KNC). **(e–f)** The same for Spearman correlation between pairwise distances (CPD).

To demonstrate that our approach can also handle UMI-based transcriptomic data, we considered three further data sets. First, we analysed a *n* = 44 808 mouse retina data set from Macosko et al. (2015). Our t-SNE result preserved much of the global geometry (Figure 4a): e.g. multiple amacrine cell clusters (green), bipolar cell clusters (blue), and non-neural clusters (magenta) were placed close together. The t-SNE analysis performed by the authors in the original publication relied on downsampling and had a worse representation of the cluster hierarchy.

**Figure 4:**
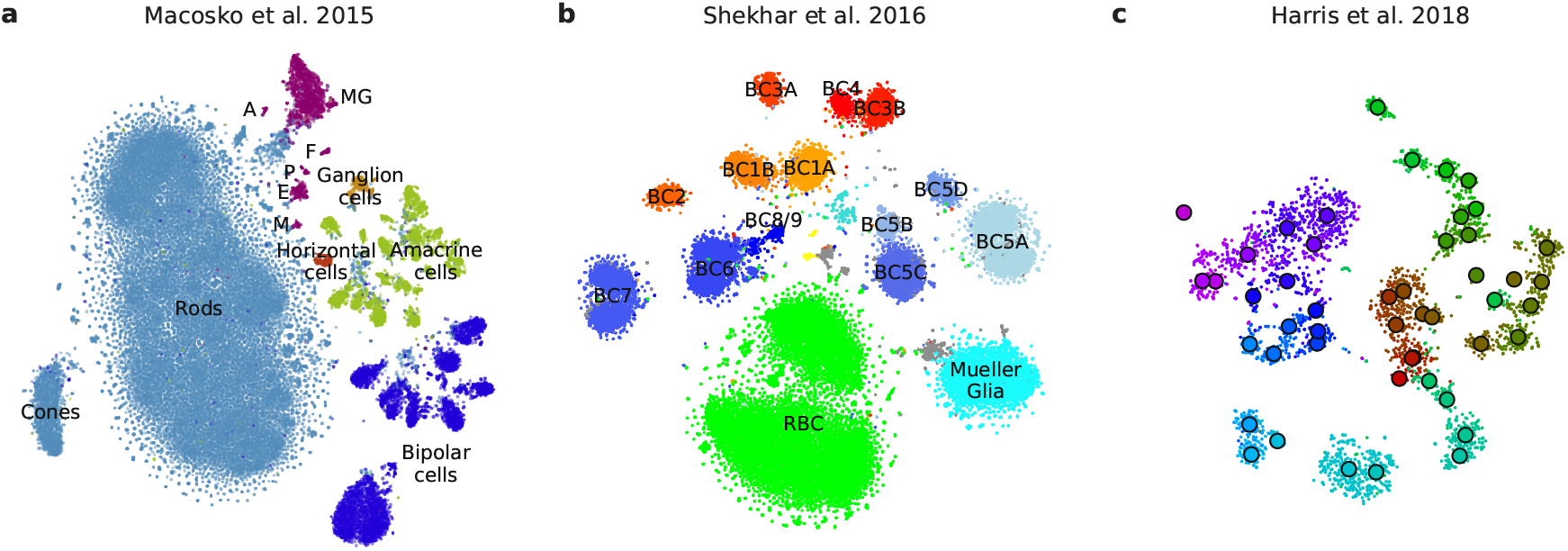
UMI-based data sets. Cluster assignments and cluster colours are taken from the original publications. **(a)** Macosko et al. (2015) data set, *n* = 44 808 cells from the mouse retina. Bipolar cells comprise eight clusters, amacrine cells comprise 21 clusters. Non-neural clusters are abbreviated (MG: Mueller glia, A: astrocytes, F: fibroblasts, P: pericytes, E: endothelium, M: microglia). **(b)** Shekhar et al. (2016) data set, *n* = 27 499 cells from the mouse retina, mostly bipolar cells. BC: bipolar cell, RBC: rod bipolar cell. Putative doublets/contaminants shown in grey. Yellow: rod and cone photoreceptors; cyan: amacrine cells. Some clusters appear to consist of two parts; this is due to an experimental batch effect that we did not remove. **(c)** Harris et al. (2018) data set, *n* = 3 663 hippocampal interneurons. Circles denote cluster centroids. The centroid of one cluster (*Sst Cryab*) is not shown because its cells were scattered all over the embedding (as they did in the original publication as well). Cluster labels not shown for visual clarity.

Second, we analysed a *n* = 27 499 data set from Shekhar et al. (2016) who sequenced cells from mouse retina targeting bipolar neurons. Here again, our t-SNE result (Figure 4b) is consistent with the global structure of the data: for example, OFF bipolar cells (types 1 to 4, warm colours) and ON bipolar cells (types 5 to 9, cold colours) are located close to each other, and four subtypes of type 5 are also close together. This was not true for the t-SNE shown in the original publication. This data set shows one limitation of our method: the data contain several very distinct but very rare clusters and those appear in the middle of the embedding, instead of being placed far out on the periphery (see Discussion).

Finally, we analysed a *n* = 3 663 data set of hippocampal interneurons from Harris et al. (2018). The original publication introduced a novel clustering and feature selection method based on the negative binomial distribution, and used a modified negative binomial t-SNE procedure. Our t-SNE visualisation (Figure 4c) did not use any of that but nevertheless led to an embedding very similar to the one shown in the original paper. Note that for data sets of this size, our method uses perplexity and learning rate that are close to the default ones.

### 2.3 Positioning new points on an existing t-SNE atlas

A common task in single-cell transcriptomics is to match a given cell to an existing reference data set. For example, introducing a protocol called Patch-seq, Cadwell et al. (2016) performed patch-clamp electrophysiological recordings followed by RNA sequencing of inhibitory cells in layer 1 of mouse visual cortex. Given the existence of the much larger Tasic et al. data set described above, it is natural to ask where on the Figure 2f, taken as a reference atlas, Cadwell et al. cells should be positioned.

It is often claimed that t-SNE does not allow out-of-sample mapping, i.e. no new points can be put on a t-SNE atlas after it is constructed. What this means is that t-SNE is a nonparametric method that does not construct any mapping *f*(x) from a high-dimensional to the low-dimensional space (parametric t-SNE is possible, but is out of scope of this paper, see Discussion). However, despite the absence of such an *f*(x), there is a straightforward way to position a new x on an existing t-SNE atlas (e.g. Berman et al., 2014; Macosko et al., 2015).

It can be done as follows: for each Cadwell et al. cell (*n* = 46), we found its *k* = 10 nearest neighbours among the Tasic et al. reference cells, using Pearson correlation across the most variable Tasic et al. genes as distance (Kiselev et al., 2018). Then we positioned the cell at the median t-SNE location of these *k* reference cells (Figure 5a). The result agreed very well with the assignment of Cadwell et al. cells to the Tasic et al. clusters performed in Tasic et al. (2018).

**Figure 5:**
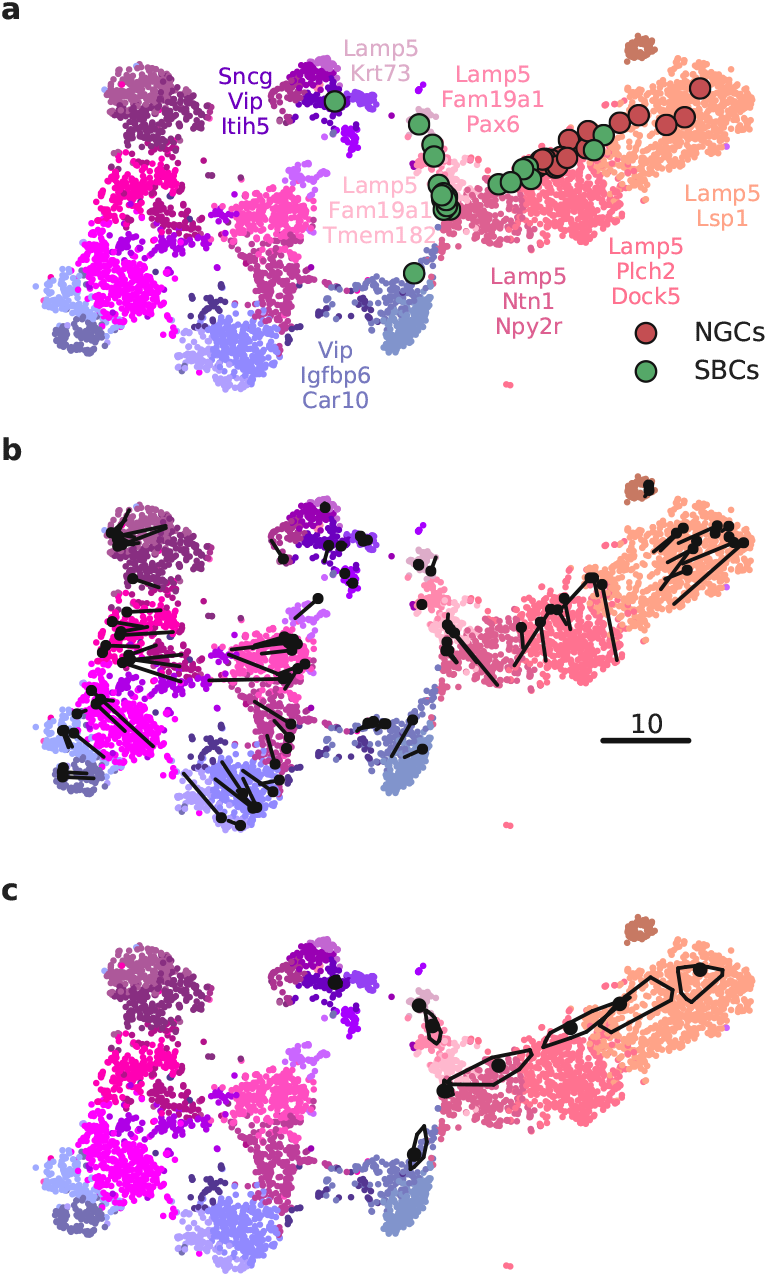
Out-of-sample mapping. **(a)** Interneurons from Cadwell et al. (2016) positioned on the Tasic et al. (2018) reference t-SNE atlas from Figure 2f. Only *Vip*/*Lamp5* continent from Figure 2f is shown here, as no cells mapped elsewhere. Cluster labels are given only for clusters where at least one cell mapped to. NGC: neurogliaform cells; SBC: single bouquet cells. Two cells out of 46 are not shown because they had an ambiguous class assignment. **(b)** A cross-validation approach: 100 random *Vip*/*Lamp5* cells from the Tasic et al. data set were removed from the t-SNE embedding and then positioned back on it using our method. Black dots show new positions, black lines connect them to the original t-SNE locations of the same cells. **(c)** Positioning uncertainty for several exemplary cells from panel (a). Polygons are convex hulls covering 95% of bootstrap repetitions.

An important caveat is that this method assumes that for each new cell there are cells of the same type in the reference data set. Cells that do not have a good match in the reference data can end up positioned in a misleading way. However, this assumption is justified whenever cells are mapped to a comprehensive reference atlas covering the same tissue, as in the example case shown here.

In a more sophisticated approach (Berman et al., 2014; Macosko et al., 2015), each new cell is initially positioned as outlined above but then its position is optimised using the t-SNE loss function: the cell is attracted to its nearest neighbours in the reference set, with the effective number of nearest neighbours determined by the perplexity. We found that the simpler procedure without this additional optimisation step worked well for our data; the additional optimisation usually has only a minor effect.

We can demonstrate the consistency of our method by a procedure similar to a leave-one-out crossvalidation. We repeatedly removed one random Tasic et al. cell from the *Vip*/*Lamp5* clusters, and positioned it back on the same reference t-SNE atlas (excluding the same cell from the *k* = 10 nearest neighbours). Across 100 repetitions, the average distance between the original cell location and the test positioning was 3.2 ± 2.4 (mean±SD; see Figure 5b for a scale bar), and most test cells stayed in the same cluster.

Positioning uncertainty can be estimated using bootstrapping across genes (inspired by Tasic et al., 2018). For each of the Cadwell et al. cells, we repeatedly selected a bootstrapped sample from the set of highly variable genes and repeated the positioning procedure (100 times). This yielded a set of bootstrapped mapping locations; the larger the variability in this set, the larger the uncertainty. To visualise the uncertainty, we show a convex hull covering 95% of the bootstrap repetitions (Figure 5c), which can be interpreted as a confidence ‘interval’. A large polygon means high uncertainty; a small polygon means high precision. For some cells the polygons are so small that they are barely visible in Figure 5c. For some other cells the polygons are larger and sometimes spread across the border of two adjacent clusters. This suggests that the cluster assignments for these cells are not certain.

### 2.4 Aligning two t-SNE visualisations

Tasic et al. (2018) is a follow-up to Tasic et al. (2016) where *n* = 1 679 cells from mouse visual cortex were sequenced with an earlier sequencing protocol. If one excludes from the new data set all clusters that have cells mostly from outside of the visual cortex, then the remaining data set has *n* = 19 366 cells. How similar is the cluster structure of this newer and larger data set compared to the older and smaller one? One way to approach this question is through aligned t-SNE visualisations.

To obtain an aligned t-SNE visualisation, we made t-SNE visualisation of the Tasic et al. (2016) data set using PCA initialisation and perplexity 50 (Figure 6a). We positioned cells of the Tasic et al. (2018) data set on this reference using the procedure described above and used the resulting layout as initialisation for t-SNE (with learning rate *n*/12 and perplexity combination of 30 and *n*/100, as elsewhere). The resulting t-SNE embedding is aligned to the previous one (Figure 6b). Of course the larger data set has higher resolution and was clustered into more clusters. But the paired and aligned t-SNE visualisations allow to make more detailed comparisons.

**Figure 6:**
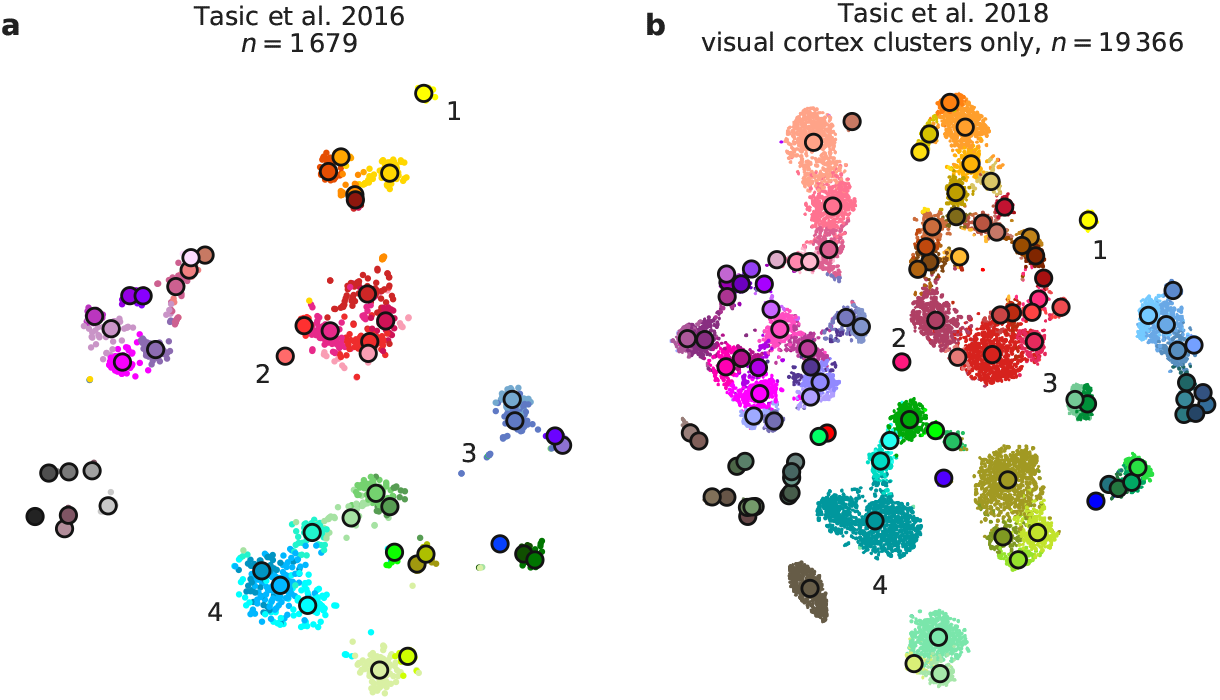
Aligned embeddings. **(a)** T-SNE visualisation of the Tasic et al. (2016) data set. Cluster assignments and cluster colours are taken from the original publication. Circles show cluster centroids. Numbers highlight some noteworthy cases, see text. **(b)** T-SNE visualisation of the Tasic et al. (2018) data set after excluding all clusters that mostly consisted of cells from anterior lateral motor cortex (23 clusters that have “ALM” in the cluster name). This t-SNE analysis was initialised by positioning all cells on the reference atlas from panel (a), ensuring that the two panels are aligned with each other.

Several noteworthy observations are highlighted in Figure 6. (1) and (2) are examples of well-isolated clusters in the 2016 data that remained well-isolated in the 2018 data (*Sst Chodl* and *Pvalb Vipr2*; here and below we use the 2018 nomenclature). (3) is an example of a small group of cells that was not assigned to a separate cluster back in 2016, became separate on the basis of the 2018 data, but in retrospect appears well-isolated already in the 2016 t-SNE plot (two *L5 LP VISp Trhr* clusters). Finally, (4) shows an example of several clusters in the 2016 data merging together into one cluster based on the 2018 data (*L4 IT VISp Rspo1*). These observations are in good correspondence with the conclusions of Tasic et al. (2018), but we find that t-SNE adds a valuable perspective and allows for an intuitive comparison.

### 2.5 Preserving global geometry with t-SNE: large data sets

Performing t-SNE on large data sets with *n* ≫ 100 000 presents several additional challenges to those already discussed above. First, vanilla t-SNE (van der Maaten and Hinton, 2008) is slow for *n* ≫ 1000 and computationally unfeasible for *n* ≫ 10 000. A widely used approximation called Barnes-Hut t-SNE (van der Maaten, 2014) in turn becomes very slow for *n* ≫ 100 000 (see Box 2). For larger data sets a faster approximation scheme is needed. This challenge was effectively solved by Linderman et al. (2019) who developed a novel t-SNE approximation based on an interpolation scheme accelerated by the fast Fourier transform (see Box 2), called FIt-SNE. Using FIt-SNE, we were able to process a data set with 1 million points and 50 dimensions (perplexity 30) in 29 minutes on a computer with four 3.4 GHz double-threaded cores, and in 11 minutes on a server with twenty 2.2 GHz double-threaded cores.

#### Box 2: The t-SNE optimisation

The original publication (van der Maaten and Hinton, 2008) suggested optimising 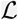 using adaptive gradient descent with momentum. They initialised **y**_*i*_ with a standard Gaussian distribution with standard deviation 0.0001. It is important that initial values have small magnitude: otherwise optimisation fails to converge to a good solution.

To escape bad local minima and allow similar points to be quickly pulled together, van der Maaten and Hinton (2008) suggested an ‘early exaggeration’ trick: during initial iterations they multiplied all attractive forces by *α* > 1. Later work (van der Maaten, 2014) used *α* = 12 for the firs 250 iterations, which became the default since then.

The exact t-SNE computes *n*^2^ similarities *p_ij_* and *n*^2^ pairwise attractive and repulsive forces on each gradient descent step. This becomes unfeasible for *n* ≫ 10 000. In the follow-up paper, van der Maaten (2014) suggested two approximations in order to speed up the computations. First, he noticed that for any perplexity value 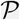 all but 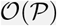 nearest neighbours of any given point *i* will have nearly zero values *p*_*j*|*i*_. He suggested to only find 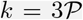 nearest neighbours of each point and set *p*_*j*|*i*_ = 0 for the remaining 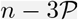 points. This relied on finding the exact 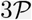 nearest neighbours, but in later work various authors (Pezzotti et al., 2017; Tang et al., 2016; Linderman et al., 2019; McInnes et al., 2018) started using approximate nearest neighbour algorithms which is much faster and does not seem to make t-SNE results any worse.

Using 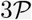 nearest neighbours accelerates computation of the attractive forces. To accelerate the repulsive force computations, van der Maaten (2014) used the Barnes-Hut approximation, originally developed for N-body simulations in physics. This reduces computational complexity from 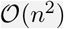 to 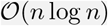, works reasonably fast for *n* up to ~100 000, but becomes too slow for much larger sample sizes. Inspired by the fast multipole method, another technique originally developed for N-body simulations, Linderman et al. (2019) suggested to interpolate repulsive forces on an equispaced grid and to use fast Fourier transform to accelerate the interpolation (FIt-SNE). This lowers computational complexity to 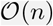 and works very fast for *n* into millions. Throughout the paper, we used their C++ implementation available at https://github.com/klugerlab/FIt-SNE.

Second, for *n* ≫ 100 000, t-SNE with default optimisation parameters tends to produce poorly converged solutions and embeddings with continuous clusters fragmented into several parts. Various groups (e.g. Wolf et al., 2018; Linderman et al., 2019) have noticed that these problems can be alleviated by increasing the number of iterations, the length or strength of the early exaggeration (see Box 2), or the learning rate. Belkina et al. (2018) demonstrated in a thorough investigation that dramatically increasing the learning rate from the default value *η* = 200 to *η* = *n*/12 prevents cluster fragmentation during the early exaggeration phase and yields a well-converged solution within the default 1000 iterations.

Third, for *n* ≫ 100 000, t-SNE embeddings tend to become very ‘crowded’, with little white space even between well-separated clusters (Unen et al., 2017). The exact mathematical reason for this is not fully understood, but the intuition is that the default perplexity becomes too small compared to the sample size, repulsive forces begin to dominate, and clusters blow up and coalesce like adjacent soap bubbles. While so far there is no principled solution for this in the t-SNE framework, a very practical trick suggested by Linderman et al. (2017) is to increase the strength of all attractive forces by a small constant ‘exaggeration’ factor between 1 and ~10 (see Methods). This counteracts the expansion of the clusters.

Fourth, our approach to preserve global geometry relies on using large perplexity *n*/100 and becomes computationally unfeasible for *n* ≫ 100 000 because FIt-SNE runtime grows linearly with perplexity. For such sample sizes, the only practical possibility is to use perplexity values in the standard range 10–100. To address this challenge, we make an assumption that global geometry should be detectable even after strong downsampling of the data set. This suggests the following pipeline: (i) downsample a large data set to some manageable size; (ii) run t-SNE on the subsample using our approach to preserve global geometry; (iii) position all the remaining points on the resulting t-SNE plot using nearest neighbours; (iv) use the result as initialisation to run t-SNE on the whole data set.

We demonstrate these ideas using two currently largest scRNA-seq data sets. The first one is a 10x Genomics data set with *n* = 1 306 127 cells from mouse embryonic brain. We first created a t-SNE embedding of a randomly selected subset of *n* = 25 000 cells (Figure 7a). As above, we used PCA initialisation, perplexity combination of 30 and *n*/100 = 250, and learning rate *n*/12. We then positioned all the remaining cells on this t-SNE embedding using their nearest neighbours (here we used Euclidean distance in the PCA space, and *k* = 10 as above; this takes ~10 minutes). Finally, we used the result as initialisation to run t-SNE on all points using perplexity 30, exaggeration coefficient 4, and learning rate *n*/12 (Figure 7b).

**Figure 7:**
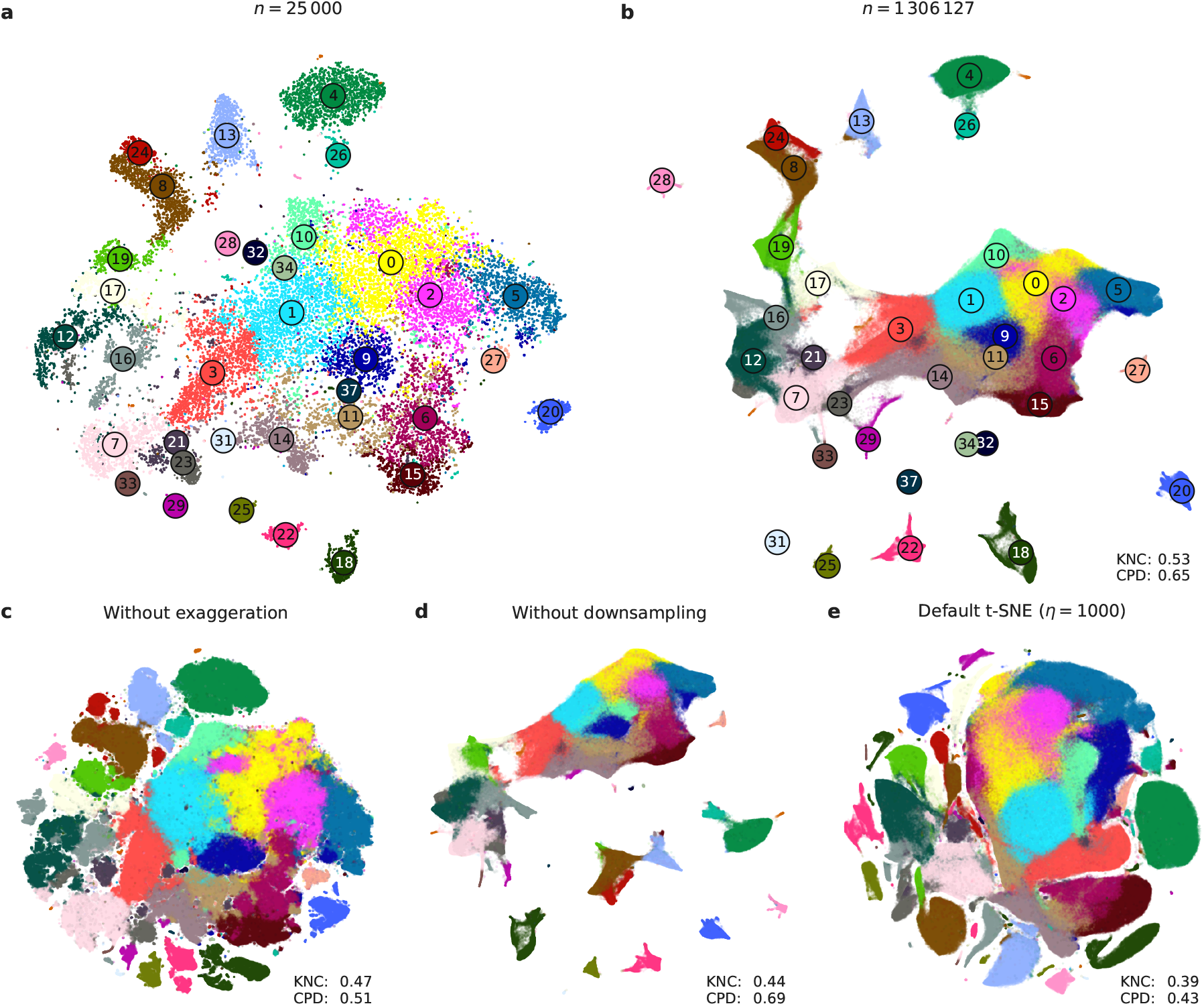
10x Genomics data set. Sample size *n* = 1 306 127. Cluster assignments and cluster colours are taken from Wolf et al. (2018). **(a)** T-SNE of a random subsample of 25 000 cells (PCA initialisation, perplexity combination of 30 and 250, learning rate 25 000/12). Cluster labels for several small clusters (30, 35, 36, and 38) are not shown here and in (b) because these clusters were very dispersed in the embeddings. **(b)** T-SNE of the full data set. All cells were positioned on the embedding in panel (a) and this was used as initialisation. Perplexity 30, exaggeration 4, learning rate *n*/12. **(c)** The same as in (b) but without exaggeration. **(d)** The same as in (b) but with PCA initialisation, i.e. without using the downsampling step. **(e)** Default t-SNE with learning rate set to *η* = 1000: random initialisation, no exaggeration.

To validate our procedure, we identified meaningful biological structures in the embedding using developmental marker genes (Englund et al., 2005; Yuzwa et al., 2017; Iacono et al., 2018). The left part of the main continent is composed of radial glial cells expressing *Aldoc* and *Slc1a3* (Figure 8a). The neighbouring areas consist of neural progenitors (neuroblasts) expressing *Eomes*, previously known as *Tbr2* (Figure 8b). The right part of the main continent consists of mature excitatory neurons expressing pan-neuronal markers such as *Stmn2* and *Tubb3* (Figure 8c) but not expressing inhibitory neuron markers *Gad1* or *Gad2* (Figure 8d), whereas the upper part of the embedding is occupied by several inhibitory neuron clusters (Figure 8d). This confirms that our t-SNE embedding shows meaningful topology and is able to capture the developmental trajectories: from radial glia, to excitatory/inhibitory neural progenitors, to excitatory/inhibitory mature neurons.

**Figure 8:**
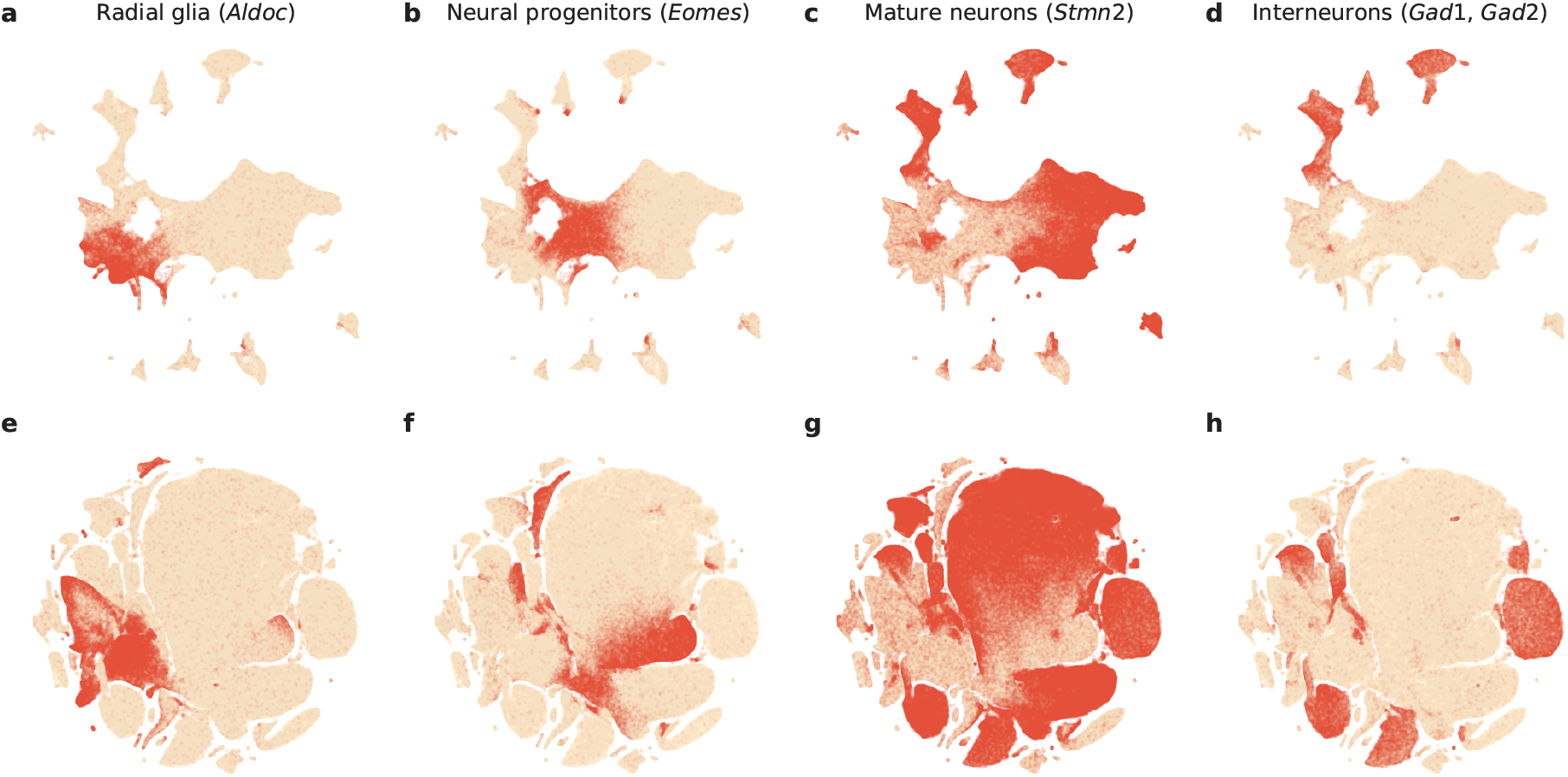
Developmental marker genes. Overlay over t-SNE embeddings from Figure 7. **(a)** Expression of *Aldoc* gene (marker of radial glia) on the t-SNE embedding from Figure 7b. Any cell with *Aldoc* detected (UMI count above zero) was coloured in red. Another radial glia marker, *Slc1a3*, had similar but a bit broader expression. **(b)** Expression of *Eomes*, marker of neural progenitors (neuroblasts). **(c)** Expression of *Stmn2*, marker of mature neurons. A pan-neuronal marker *Tubb3* had similar but a bit broader expression. **(d)** Expression of *Gad1* and *Gad2* (either of them), markers of inhibitory neurons. **(e–h)** The same genes overlayed over the default t-SNE embedding from Figure 7e.

We illustrate the importance of the components of our pipeline by a series of control experiments. Omitting exaggeration yields over-expanded clusters and a plot with less discernible global structure (Figure 7c). Without downsampling, the global geometry is preserved worse (Figure 7d): e.g. most of the interneuron clusters are in the lower part of the figure, but clusters 17 and 19 (developing interneurons), are located in the upper part. Finally, the default t-SNE with random initialisation and no exaggeration (but learning rate set to *η* = 1000) yields a poor embedding that fragments some of the clusters and misrepresents global geometry (Figure 7e). Indeed, overlaying the same marker genes shows that developmental trajectories are not preserved and related groups of cells, e.g. interneurons, are dispersed across the embedding (Figure 8e–h). Again, this is not a strawman: this embedding is qualitatively similar to the ones shown in the literature (Wolf et al., 2018; Bhaduri et al., 2018).

In addition, we analysed a data set encompassing *n* = 2 058 652 cells from mouse embryo at several developmental stages (Cao et al., 2019). The original publication showed a t-SNE embedding that we reproduce in Figure 9a. Whereas it shows a lot of structure, it visibly suffers from all the problems mentioned above: some clusters are fragmented into parts (e.g. clusters 13 and 15), there is little separation between distinct cell types, and global structure is grossly misrepresented. The authors annotated all clusters and split them into ten biologically meaningful developmental trajectories; these trajectories are intermingled in Figure 9a. In contrast, our t-SNE embedding (Figure 9b) neatly separates all ten developmental trajectories and arranges clusters within major trajectories in a meaningful developmental order: e.g. there is a continuous progression from radial glia (cluster 7), to neural progenitors (9), to postmitotic premature neurons (10), to mature excitatory (5) and inhibitory (15) neurons.

**Figure 9:**
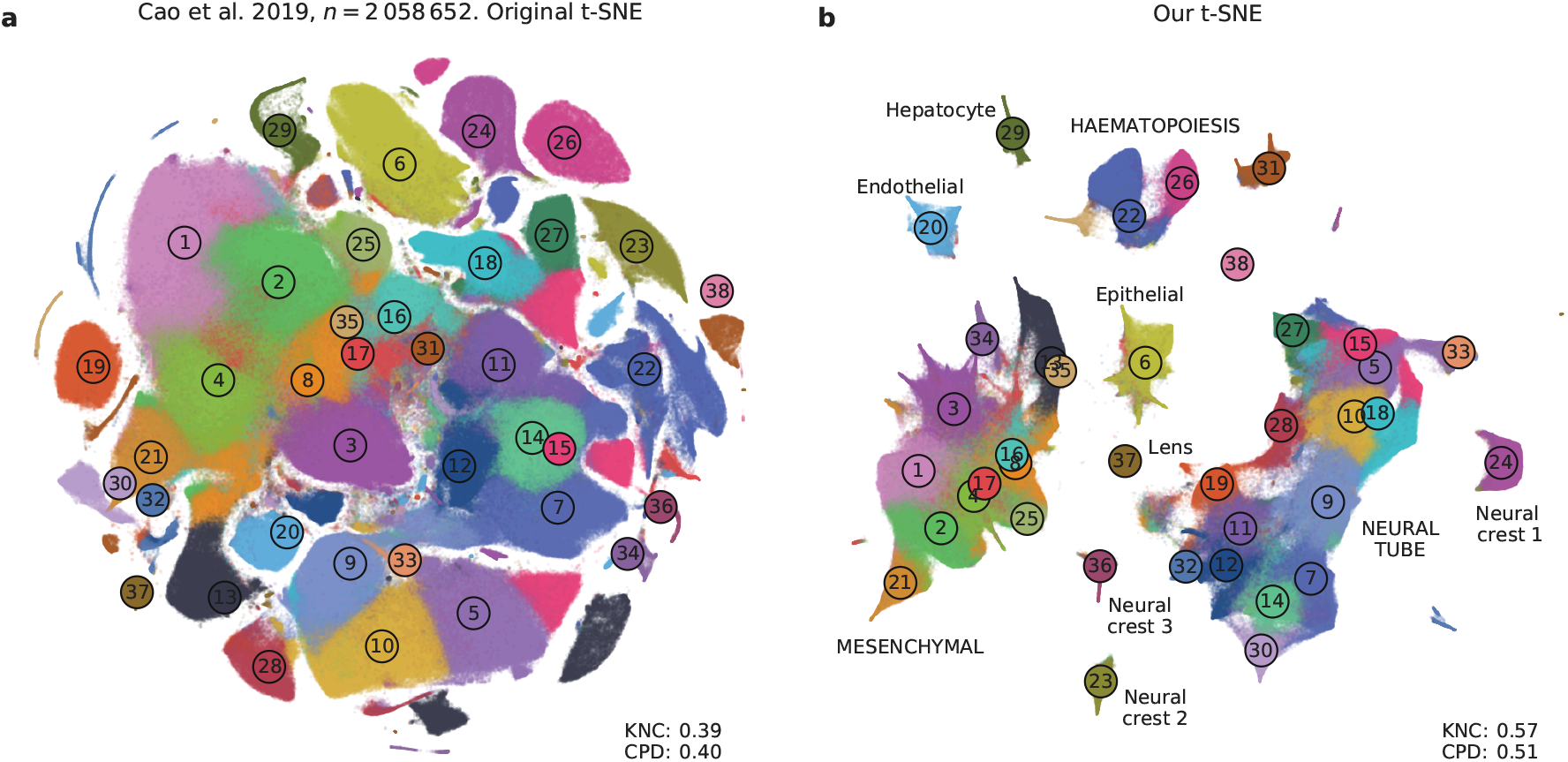
Cao et al. data set. Sample size *n* = 2 058 652. Cluster assignments and cluster colours are taken from the original publication (Cao et al., 2019). **(a)** T-SNE embedding from the original publication. The authors ran t-SNE in `scanpy` with default settings, i.e. with random initialisation, perplexity 30, and learning rate 1000. For cluster annotations, see original publication. **(b)** T-SNE embedding produced with our pipeline for large data sets: a random sample of 25 000 cells was embedded using PCA initialisation, learning rate 25 000/12, and perplexity combination of 30 and 250; all other cells were positioned on resulting embedding and this was used to initialise t-SNE with learning rate 2 058 652/12, perplexity 30, and exaggeration 4. Labels correspond to the ten developmental trajectories identified in the original publication. Labels in capital letters denote trajectories consisting of multiple clusters. 32 011 putative doublet cells are not shown in either panel.

### 2.6 Comparison with UMAP

A promising dimensionality reduction method called UMAP (McInnes et al., 2018) has recently attracted considerable attention in the transcriptomics community (Becht et al., 2019). Technically, UMAP is very similar to an earlier method called largeVis (Tang et al., 2016), but McInnes et al. (2018) provided a mathematical foundation and a convenient Python implementation. LargeVis and UMAP use the same attractive forces as t-SNE does, but change the nature of the repulsive forces and use a different, sampling-based approach to optimisation. UMAP has been claimed to be faster than t-SNE and to outperform it in terms of preserving the global structure of the data (McInnes et al., 2018; Becht et al., 2019).

While UMAP is indeed much faster than Barnes-Hut t-SNE, FIt-SNE (Linderman et al., 2019) is at least as fast as UMAP. We found FIt-SNE 1.1 with default settings to be ~4 times faster than UMAP 0.3 with default settings when analysing the 10x Genomics (14 m vs. 56 m) and the Cao et al. (31 m vs. 126 m) data sets on a server with twenty 2.2 GHz double-threaded cores (for this experiment, the input dimensionality was 50 and the output dimensionality was 2; UMAP may be more competitive in other settings). That said, the exact run time will depend on the details of implementation, and both methods may be further accelerated in future releases, or by using GPU parallelisation (Chan et al., 2019).

To compare UMAP with our t-SNE approach in terms of preservation of global structure, we first ran UMAP on the synthetic and on the Tasic et al. 2018 data sets (Figure S2). We used the default UMAP parameters, and also modified the two key parameters (number of neighbours and tightness of the embedding) to produce a more ‘t-SNE-like’ embedding. In both cases and for both data sets, all three metrics (KNN, KNC, and CPD) were considerably lower than with our t-SNE approach.

Next, we applied UMAP with default parameters to the 10x Genomics and the Cao et al. data sets (Figure S3). For the Cao et al. data set, the UMAP embedding was qualitatively similar to our t-SNE embedding, but possibly misrepresented some aspects of the global topology (Figure S3b). For the 10x Genomics data set, the UMAP embedding was also qualitatively similar to our t-SNE embedding (Figure 7b), but the global geometry again seemed worse because e.g. the interneuron clusters were scattered around the embedding (Figure S3a). Notably, we observed that the global structure of UMAP embeddings strongly depended on the random seed.

An in-depth comparison of t-SNE and UMAP is beyond the scope of our paper, but this analysis suggests that previous claims that UMAP vastly outperforms t-SNE (Becht et al., 2019) might have been partially due to t-SNE being applied in suboptimal way. Our analysis also indicates that UMAP does not necessarily solve t-SNE’s problems out of the box and might require as many careful parameter and/or initialisation choices as t-SNE does. Many recommendations for running t-SNE that we made in this manuscript can likely be adapted for UMAP.

## 3 Discussion

Here we described how to set t-SNE parameters and initialisation in order to create accurate lowdimensional visualisations of high-dimensional scRNA-seq data sets that preserve global, hierarchical structure. Accordingly, the resulting visualisations are biologically more interpretable compared to the default t-SNE settings. Our procedure works well even for large data sets and — due to the PCA initialisation — the visualisation is deterministic and does not depend on a random seed. In addition, we provide means to create aligned presentations of multiple data sets and to map additionally collected cells onto existing visualisations.

### 3.1 Preserving global structure

The fact that t-SNE does not always preserve global structure is one of its well-known limitations. As formulated in Wattenberg et al. (2016), “the basic message is that distances between well-separated clusters in a t-SNE plot may mean nothing.” Indeed, the algorithm, by construction, only cares about preserving local neighbourhoods.

We showed that using informative initialisation (such as PCA initialisation, or downsampling-based initialisation) can substantially improve the global structure of the final embedding because it ‘survives’ through the optimisation process. Importantly, a custom initialisation does not interfere with t-SNE optimisation and does not yield a worse solution (Figure 3a,b). Standard t-SNE uses a random Gaussian initialisation with very small variance (Box 2), and whenever we use PCA or downsampling-based initialisation, we always scale it to have the same small variance. As a result, the final Kullback-Leibler divergence (Box 1) after using custom (but appropriately scaled) initialisation was never higher than after using a random initialisation. Another possible concern is that a custom initialisation might bias the resulting embedding by ‘injecting’ some artefact global structure. However, if anything may be seen as ‘injecting’ artefactual structure, it is rather the *random* initialisation. Indeed, the global arrangement of clusters on a standard t-SNE embeddings often strongly depends on the random seed.

Apart from the initialisation, we suggested that using large perplexity values — substantially larger than the commonly used ones — can be useful in the scRNA-seq context. Our experiments suggest that whereas PCA initialisation helps preserving the macroscropic structure, large perplexity (either on its own or as part of a perplexity combination) helps preserving the mesoscopic structure (Figure 3d,f).

It has recently been claimed that UMAP preserves the global geometry better than t-SNE (Becht et al., 2019). However, UMAP operates on the *k*-nearest neighbour graph, exactly as t-SNE does, and is therefore not designed to preserve large distances any more than t-SNE. To give a specific example, Cao et al. (2019) performed both t-SNE and UMAP and observed that “unlike t-SNE, UMAP places related cell types near one another”. We demonstrated that this is largely because t-SNE parameters were not set appropriately. Simply using a high learning rate *n*/12 (even without any initialisation tricks) “places related cell types near one another” as well as UMAP does, and additionally using exaggeration factor 4 separates clusters in more compact groups, similar to UMAP.

### 3.2 Parameter choice for t-SNE

Often, t-SNE is perceived as having only one free parameter to tune, perplexity. Under the hood, however, there are also various optimisation parameters (such as the learning rate, the number of iterations, early exaggeration factor, etc.) and we showed above that they can have a dramatic effect on the quality of the visualisation. Here we have argued that exaggeration can be used as another useful parameter when embedding large data sets. In addition, while the low-dimensional similarity kernel in t-SNE has traditionally been fixed as the t-distribution with *ν* =1 degree of freedom, we showed in a parallel work that modifying *ν* can uncover additional fine structure in the data (Kobak et al., 2019).

One may worry that this gives a researcher ‘too many’ knobs to turn. Above, we have given clear guidelines on how to set these parameters for effective visualisations. As argued above and in Belkina et al. (2018), setting the learning rate to *n*/12 ensures good convergence and automatically takes care of the optimisation issues. Perplexity should be left at the default value 30 for very large data sets, but can be combined with *n*/100 for smaller data sets. Exaggeration can be increased to ~4 for very large data sets, but is not needed for smaller data sets.

In comparison, UMAP has two main adjustable parameters (and many further optimisation parameters): `n_neighbors`, corresponding to perplexity, and `min_dist`, controlling how tight the clusters become. The latter parameter sets the shape of the low-dimensional similarity kernel (McInnes et al., 2018) and is therefore analogous to *ν* mentioned above. Furthermore, our experiments with UMAP suggest that its repulsive forces roughly correspond to t-SNE with exaggeration ~4 (Figures S2, S3 in comparison with Figures 1, 2, 7, 9). Whether this is desirable, depends on the application. With t-SNE, one can choose to switch exaggeration off and e.g. use the embedding shown in Figure 7c instead of Figure 7e. Our pipeline will still ensure that the global structure is about right.

### 3.3 Comparison to related work

Several variants of t-SNE have been recently proposed in the literature. One important example is parametric t-SNE, where a neural network is used to create a mapping *f*(x) from high-dimensional input space to two dimensions (i.e. the output layer of the network contains two neurons) and is then trained using standard deep learning techniques to yield an optimal t-SNE result (van der Maaten, 2009). One potential advantage of this approach is that the ‘most appropriate’ perplexity does not need to grow with the sample size, as long as the mini-batch size remains constant. Parametric t-SNE has been recently applied to transcriptomic data under the names net-SNE (Cho et al., 2018) and scvis (Ding et al., 2018). The latter method combined parametric t-SNE with a variational autoencoder, and was claimed to yield more interpretable visualisations than standard t-SNE due to better preserving the global structure. Indeed, the network architecture limits the form that the mapping *f*(x) can take; this implicit constraint on the complexity of the mapping prevents similar clusters from ending up in very different parts of the resulting embedding.

Another important development is hierarchical t-SNE, or HSNE (Pezzotti et al., 2016). The key idea is to use random walks on the *k*-nearest neighbours graph of the data to find a smaller set of ‘landmarks’, which are points that can serve as representatives of the surrounding points. In the next round, the *k*-nearest neighbours graph on the level of landmarks is constructed. This operation can reduce the size of the data set by an order of magnitude, and can be repeated until the data set becomes small enough to be analysed with t-SNE. Each level of the landmarks hierarchy can be explored separately. Unen et al. (2017) successfully applied this method to mass cytometry data sets with up to 15 million cells. However, HSNE does not allow to embed all *n* points in a way that would preserve the geometry on the level of landmarks.

Another related method, UMAP (McInnes et al., 2018), has been discussed above. All of these methods (net-SNE, scvis, HSNE, UMAP) can potentially become important tools for transcriptomic data analysis. Our goal in this manuscript was not to argue that t-SNE is better or worse than any of them, but to demonstrate what t-SNE itself is capable of when applied with care.

### 3.4 Limitations and future work

Our approach to preserving global geometry of the data is based on using PCA initialisation and large perplexity values. It can fail if some aspects of the global geometry are not adequately captured in the first two PCs or by the similarities computed using a large perplexity. One scenario when this is likely to happen is when the data set contains very isolated and very rare cell types. Indeed, a small isolated cluster might not appear isolated in the first two PCs because it would not have enough cells to contribute much to the explained variance. At the same time, large perplexity will make the points in that cluster be attracted to some arbitrary, unrelated, clusters. As a result, a small cluster can get ‘sucked into’ the middle of the embedding even if it is initialised on the periphery.

Among the data sets analysed in this manuscript, this happened in the Shekhar et al. data set (Figure 4b): cone and rod photoreceptor (yellow) and amacrine cell (cyan) clusters ended up in the middle of the embedding despite being very different from all the bipolar cell clusters. One simple visualisation that can show this is the MDS embedding of the cluster means (the means are unaffected by the relative abundances of the clusters); thus, when t-SNE is done together with clustering, we recommend to supplement a t-SNE visualisation with a MDS visualisation of cluster means (as in Figure 2a). Alternatively, one could use PAGA (Wolf et al., 2019), a recent method specifically designed to visualise the relationships between clusters in scRNA-seq data.

This example highlights that our approach is not a final solution to preserving the global structure of the data. A principled approach would incorporate some terms ensuring adequate global geometry directly into the loss function, while making sure that the resulting algorithm is scalable to millions of points. We consider it a very important direction for future work. In the meantime, we believe that our recommendations will strongly improve t-SNE visualisations used in the current single-cell transcriptomic studies, and may be useful in other application domains as well.

## 4 Methods

### 4.1 Code availability and tutorial code

We prepared a self-contained Jupyter notebook in Python that demonstrates all the techniques presented in this manuscript. It is available at https://github.com/berenslab/rna-seq-tsne. For simplicity, it uses the Tasic et al. (2018) data set for all demonstrations. To show how to map new cells to the reference t-SNE atlas, we split the data into a training set and a test set. To show how to align two t-SNE visualisations, we split the data set into two parts. To show how to process a much larger data set, we replicate each cell 10 times and add noise.

The code that does the analysis and produces all the figures used in this manuscript is available in form of Python notebooks at https://github.com/berenslab/rna-seq-tsne. We used a C++ implementation of FIt-SNE (Linderman et al., 2019), version 1.1, available at https://github.com/klugerlab/FIt-SNE together with interfaces for R, Matlab, and Python. While working on this paper, we contributed to this package several additional features that were crucial for our pipeline.

FIt-SNE has been re-implemented as a pure Python package `openTSNE`, available at https://github.com/pavlin-policar/openTSNE. It supports all the features used in this manuscript (and conveniently allows to position out-of-sample cells, with or without optimisation).

### 4.2 Pre-processing

All data sets were downloaded as tables of UMI or read counts. Let **X** be a *n* × *p* matrix of gene counts, with *n* and *p* being the number of cells and the number of genes respectively. We assume that zero columns (if any) have been removed. We used a standard pre-processing pipeline consisting of the following steps: (i) library size normalisation; (ii) feature selection; (iii) log-transformation; (iv) PCA. Specifically, we normalised the read counts to counts per million (CPM) and UMI counts to counts per median library size, selected 1000–3000 most variable genes using dropout-based feature selection similar to the one suggested in Andrews and Hemberg (2018), applied log_2_(*x* + 1) transform, and finally did PCA retaining the 50 leading PCs. We experimented with modifying and omitting any of these steps. Our experiments showed that log-transformation (or a similar nonlinear transformation) and feature selection are the two most important steps for adequate results. PCA mainly improves computational efficiency as it reduces the dimensionality and size of the data set before running t-SNE.

#### Library size normalisation

We normalised the read counts of each cell by the cell’s library size 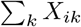, and multiplied by 1 million, to obtain counts per million (CPM). We normalised the UMI counts by the cell’s library size and multiplied by the median library size across all *n* cells in the data set (Zheng et al., 2017). This is more appropriate for UMI counts because multiplying by 1 million can strongly distort the data after the subsequent log(1 + *x*) transform (Townes et al., 2019).

#### Feature selection

Most studies use the mean-variance relationship to perform feature selection (e.g. Zheng et al., 2017): they select genes that have large variance given their mean. We adopt the approach of Andrews and Hemberg (2018) who exploit the mean-dropout relationship: the idea is to select genes that have high dropout (i.e. zero count) frequency given their mean expression level. Any gene that has high dropout rate and high mean expression could potentially be a marker of some particular subpopulation. We found it more intuitive to use the mean across non-zero counts instead of the overall mean, because it is computed independently of the fraction of zero counts.

For each gene *g*, we computed the fraction of ‘near-zero’ counts

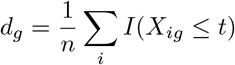

and the mean log ‘non-zero’ expression

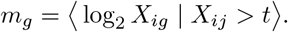

For all UMI-based data sets we used *t* = 0 and for all Smart-seq2/SMARTer data sets we used *t* = 32 (some known marker genes were not getting selected with *t* = 0). We discarded all genes that have ‘non-zero’ expression in less than *n*_min_ = 10 cells. There was a strong negative relationship between *d_g_* and *m_g_* across all the remaining genes (Figure S4). In oder to select a pre-specified number *M* of genes (usually *M* = 1000 or *M* = 3000), we used a heuristic approach of finding a value *b* such that

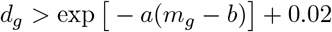

was true for exactly *M* genes. This can be done with a binary search. In Figure S4 this corresponds to moving the red line horizontally until there are exactly *M* genes to the upper-right of it. These *M* genes were then selected. We used *a* = 1.5 for all data sets apart from Macosko et al. where *a* = 1 provided a better fit for the distribution.

We performed feature selection using the raw counts (before library size normalisation). Then we used normalised values for the selected *M* genes. We used *M* = 3000 for the Tasic et al. 2018 and for the Macosko et al. data sets, and *M* = 1000 for the remaining data sets. This feature selection method was not used for the 10x Genomics and the Cao et al. data sets, see below.

#### Nonlinear transformation

We transformed all values in the *n*×*M* count matrix after feature selection with a log_2_(*x* + 1) transformation. This transformation is standard in the transcriptomics literature. It is convenient because all zeros remain zeros, and at the same time the expression counts of all genes become roughly comparable. Without this transformation, the Euclidean distances between cells are dominated by a handful of genes with very high counts. There are other transformations that can perform similarly well. In the cytometric literature, the inverse hyperbolic sine 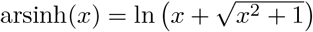 is often used, sometimes as arsinh(*x*/*r*) with *r* = 5 or a similar value (Amir et al., 2013). Note that arsinh(*x*/*r*) is the variance-stabilising transformation for the negative binomial distribution with parameter *r*, which is often taken to model UMI counts well.

#### Standardisation

Many studies standardise the *n* × *M* matrix after the log-transformation, i.e. center and scale all columns to have zero mean and unit variance. We prefer not to do this by default. In general, standardisation is recommended when different features are on different scale, but the log-transformed counts of different genes are arguably already on the same scale.

From a more theoretical point of view, if one could assume that the expression counts of each gene for cells of the same type are distributed log-normally, then Euclidean distance after log-transformation would exactly correspond to the log-likelihood. For the UMI-based data, the common assumption is that the expression counts are distributed according to the negative binomial distribution. For large counts, the negative binomial distribution behaves qualitatively similarly to the log-normal (for example, its variance function is *μ* + *μ*^2^/*r* whereas the log-normal has variance function proportional to *μ*^2^), so the Euclidean distance after the log-transformation can be thought of as approximating the negative binomial log-likelihood (Harris et al., 2018). Standardising all the genes after log-transformation would destroy this relationship.

At the same time, in some data sets we observed a stronger separation between some of the clusters after performing the standardisation step. Here we applied standardisation for those data sets in which it was used by the original authors (Macosko et al., Shekhar et al., 10x Genomics, Cao et al.).

#### Scanpy preprocessing

For the 10x Genomics and the Cao et al. data sets, we used the preprocessing pipeline `recipe_zheng17()` from `scanpy` (Wolf et al., 2018) to ease the comparison with clustering and dimensionality reduction performed by Wolf et al. (2018) and by Cao et al. (2019). This pipeline follows Zheng et al. (2017) and is similar to ours: it performs library size normalisation to median library size, selects most variable genes based on the mean-variance relationship, applies the log_2_(*x* + 1) transform and standardises each feature. We used *M* = 1000 genes for the 10x Genomics data set and *M* = 2000 for the Cao et al. data set, following the original publication.

#### Principal component analysis

We used PCA to reduce the size of the data matrix from *n* × *M* to *n* × 50 prior to running t-SNE. In our experiments, this did not have much influence on the t-SNE results but is computationally convenient. Some studies estimate the number of ‘significant’ principal components via shuffling (e.g. Shekhar et al., 2016). In our experiments, for the data sets with tens of thousands of cells, the number of significant PCs is usually close to 50 (for example, for the Tasic et al. (2018) data set it is 37, according to the shuffling test). Given that PCA does not have much influence on the t-SNE results, we prefer to use a fixed value of 50.

### 4.3 Details of the t-SNE analysis

#### Default parameters for t-SNE optimisation

Unless explicitly stated, we used the default parameters of FIt-SNE. Following (van der Maaten, 2014), the defaults are 1000 iterations with learning rate *η* = 200; momentum .5 for the first 250 iterations and .8 afterwards; early exaggeration *α* = 12 for the first 250 iterations; initialisation of the points in the 2D embedding space with coordinates drawn from a standard Gaussian distribution with standard deviation 0.0001. Further input parameters for the nearest neighbour search using the `Annoy` library (number of trees: 50, number of query nodes: 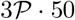 for perplexity 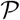) and for the grid interpolation (number of approximating polynomials: 3, maximum grid spacing: 1, minimum grid size: 50) were always left at the default values. For reproducibility, we always used random seed 42.

#### Initialisation

For PCA initialisation, we divide the first two principal components by the standard deviation of PC1 and multiply them by 0.0001, which is the default standard deviation of the random initialisation. This scaling is important: values used for initialisation should be close to zero, otherwise the algorithm might have problems with convergence (we learned about the importance of scaling from James Melville’s notes at https://jlmelville.github.io/smallvis/init.html). The same scaling was used for the downsampling-based initialisation and also for the custom initialisation when creating aligned visualisations.

The sign of the principal components is arbitrary. To increase reproducibility of the figures, we always fixed the sign of the first two PCs such that for each PCA eigenvector the sum of its values was positive (i.e. if it was negative, we flipped the sign).

#### Postprocessing

For the t-SNE embeddings of the 10x Genomics data set, we rotated the result 90 degrees clockwise and flipped horizontally, to make it visually more pleasing. Note that t-SNE result can be arbitrarily rotated and flipped as this does not change the distances between points. Caveat: it should never be stretched horizontally or vertically.

#### Multi-scale similarities

We follow Lee et al. (2015) in their definition of multi-scale similarities. For example, to combine perplexities 30 and 200, the values *p*_*j*|*i*_ (see Box 1) are computed with perplexity 30 and with perplexity 200 for each cell *i* and then averaged. This is approximately (but not exactly!) equivalent to using a different similarity kernel: instead of the Gaussian kernel 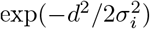 where *d* is Euclidean distance, a multi-scale kernel

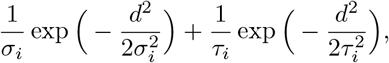

with the variances 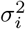 and 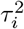 selected such that the perplexity of the first Gaussian component is 50 and the perplexity of the second Gaussian component is 500.

#### Exaggeration

Early exaggeration (see Box 2) means multiplying the attractive term in the loss function (see last equation in Box 1) by a constant *α* > 1 during the initial iterations (the default is *α* = 12 for 250 iterations). Linderman et al. (2017) suggested to use ‘late exaggeration’: using some *α* > 1 for some number of last iterations. Thus, their approach used three stages: early exaggeration, followed by no exaggeration, followed by late exaggeration. We prefer to use two stages only: we keep *α* constant after the early exaggeration is turned off. This is why we call it ‘exaggeration’ and not ‘late exaggeration’. While early exaggeration is essentially an optimisation trick (van der Maaten and Hinton, 2008), we consider subsequent exaggeration to be a meaningful change to the loss function.

#### Positioning out-of-sample cells

To position new cells on an existing t-SNE embedding we used the same *M* most variable genes that were used to create the target embedding. Usually only a subset of *L* < *M* genes was present in the count table of the new data set; we used all *L* genes in this subset to compute the correlation distances. We used coordinate-wise median among the *k* nearest neighbours.

We used correlation distances as possibly more robust for batch effects than Euclidean distances: when out-of-sample cells were sequenced with a different protocol, the batch effect can be very strong. This consideration does not apply to positioning cells for a downsampling-based initialisation (there is no batch effect by construction). For computational simplicity, we used Euclidean distance in the space of the 50 PCs in this case.

#### Bootstrapping over genes

We used bootstrapping to estimate the uncertainty of the mapping of new cells to the existing t-SNE visualisation. Given a set of *L* genes that are used for mapping, we selected a bootstrap sample of *L* genes out of *L* with repetition and performed the positioning procedure using this sample of genes. This constitutes one bootstrap iteration. We did 100 iterations and, for each cell, obtained 100 positions on the t-SNE atlas. The larger the spread of these positions, the larger the mapping uncertainty.

For Figure 5c, we computed the distances from the original mapping position to the 100 bootstrapped positions and discarded 5 bootstrap positions with the largest distance. Then we drew a convex hull of the remaining 95 bootstrap positions (using `scipy.spatial.ConvexHull`).

## Acknowledgements

We thank Anna Belkina, George Linderman, Leland McInnes, James Melville, Pavlin Poličar, and Alexander Wolf for discussions of t-SNE and UMAP. We thank Andreas Tolias for discussing patch-seq data. This work was supported by the Deutsche Forschungsgemeinschaft (BE5601/4-1 and the Cluster of Excellence “Machine Learning — New Perspectives for Science, EXC 2064, project number 390727645), the Federal Ministry of Education and Research (FKZ 01GQ1601 and 01IS18039A) and the National Institute Of Mental Health of the National Institutes of Health under Award Number U19MH114830. The content is solely the responsibility of the authors and does not necessarily represent the official views of the National Institutes of Health.

## Author contributions

DK and PB conceptualized the project, DK performed the analysis, DK and PB wrote the paper.

## Competing interests

The authors declare no competing interests.

## Data and code availability

All data were downloaded in form of count tables following links in the original publications. Full analysis scripts in Python (including direct download links) are available at https://github.com/berenslab/rna-seq-tsne.

## Supplementary Figures

**Figure S1:**
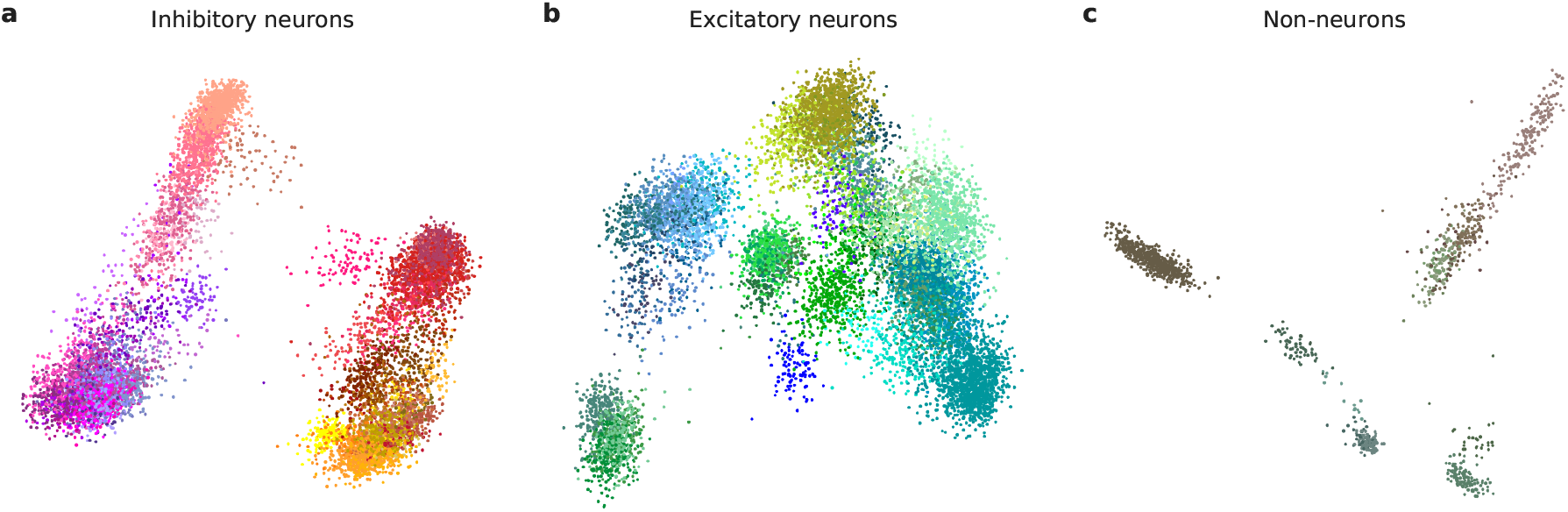
Major cell classes in the Tasic et al. data set. Cluster colours as in Figure 2. In each case PCA makes some of the within-class internal structure obvious. **(a)** PCA of the inhibitory neurons. The scatter plot shows two main principal components. Purple: *Vip* neurons. Salmon: *Lamp5* neurons. Red: *Pvalb* neurons. Orange: *Sst* neurons. **(b)** PCA of the excitatory neurons. **(c)** PCA of the non-neural cells.

**Figure S2:**
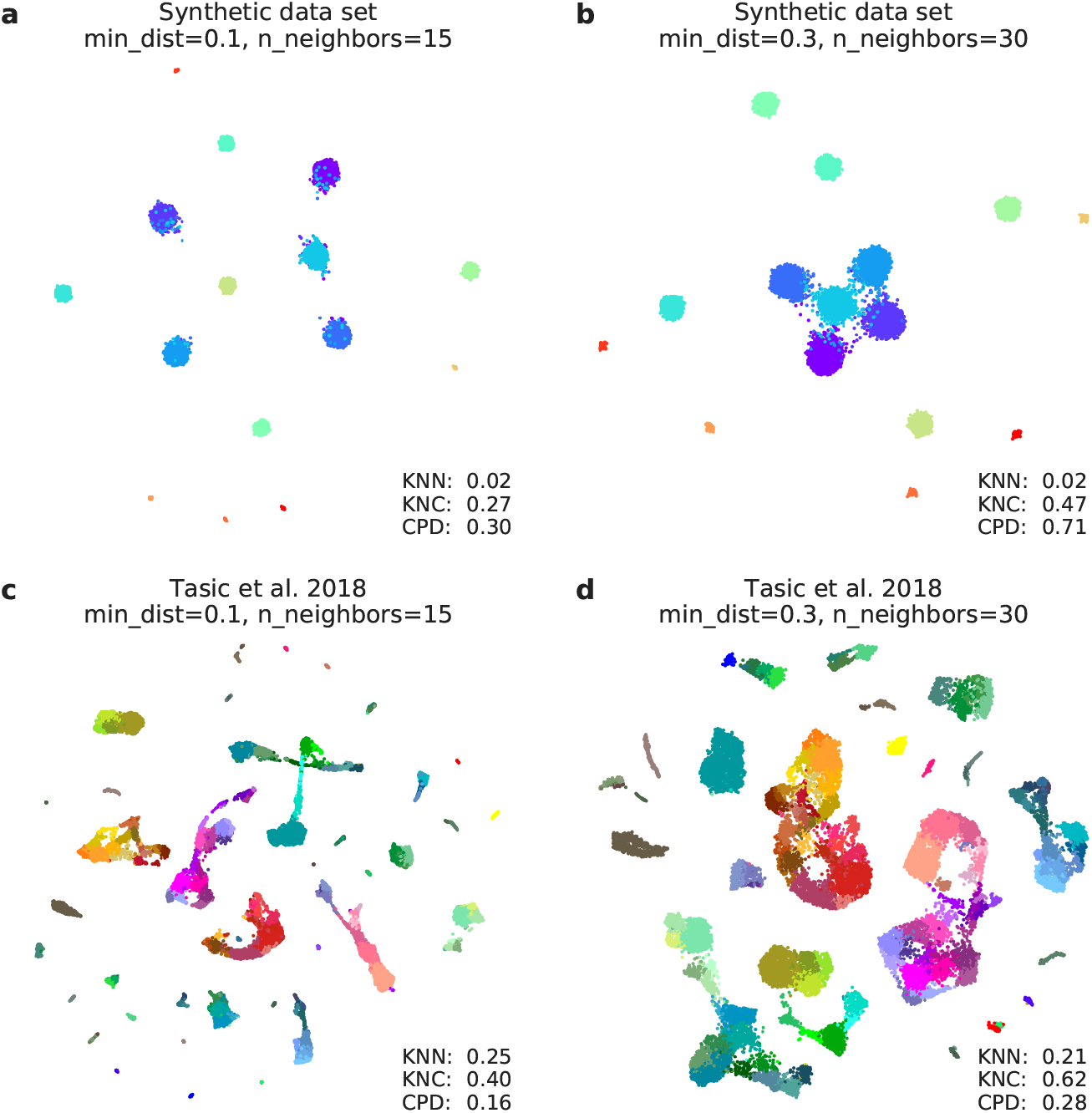
UMAP embeddings of the small data sets. Cluster colours as in Figure 1 and Figure 2. For all embeddings, random seed was set to 1. **(a)** UMAP embedding of the synthetic data set with the default parameters (`min_dist=0.1, n_neighbors=15`). **(b)** UMAP embedding of the synthetic data set with parameters `min_dist=0.5, n_neighbors=30` that tend to produce less fragmented embeddings. In both (a) and (b) all three metrics are worse than in our t-SNE embedding in Figure 1f: KNN=0.11, KNC=0.82, CPD=0.74. **(c)** UMAP embedding of the Tasic et al. 2018 data set with the default parameters (`min_dist=0.1, n_neighbors=15`). **(b)** UMAP embedding of the Tasic et al. 2018 data set with parameters `min_dist=0.5, n_neighbors=30`. In both (c) and (d) all three metrics are worse than in our t-SNE embedding in Figure 2f: KNN=0.41, KNC=0.68, CPD=0.53.

**Figure S3:**
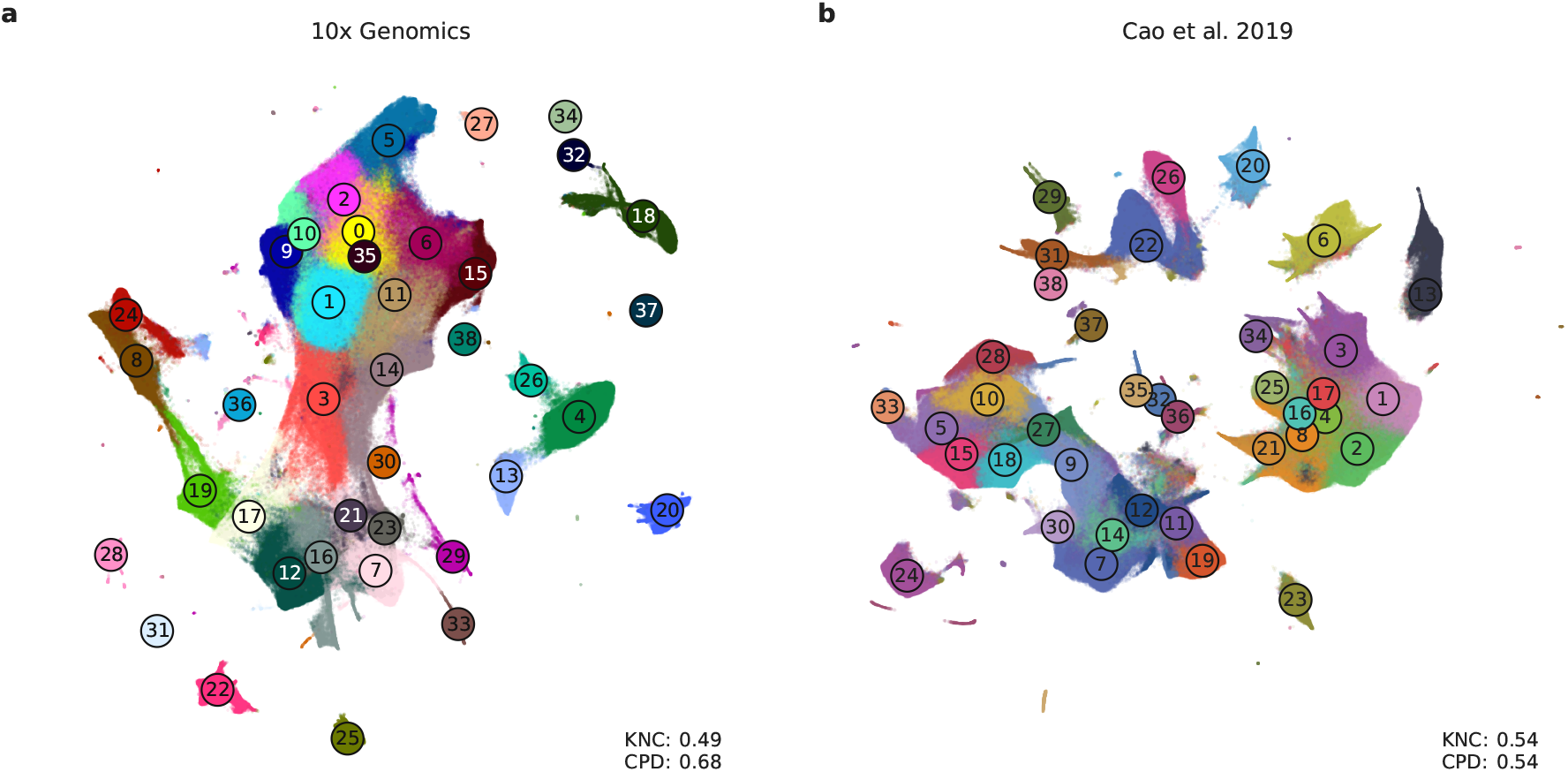
UMAP embeddings of the large data sets. Cluster colours as in Figure 7 and Figure 9. For all embeddings, random seed was set to 1. **(a)** UMAP embedding of the 10x Genomics data set, default parameters. The interneuron clusters (##8, 24, 19, 26, 4, 13) are scattered across the embedding. (b) UMAP embedding of the Cao et al. data set, default parameters. Note that cluster #13 (myocytes) appears isolated, even though it belongs to the mesenchymal trajectory.

**Figure S4:**
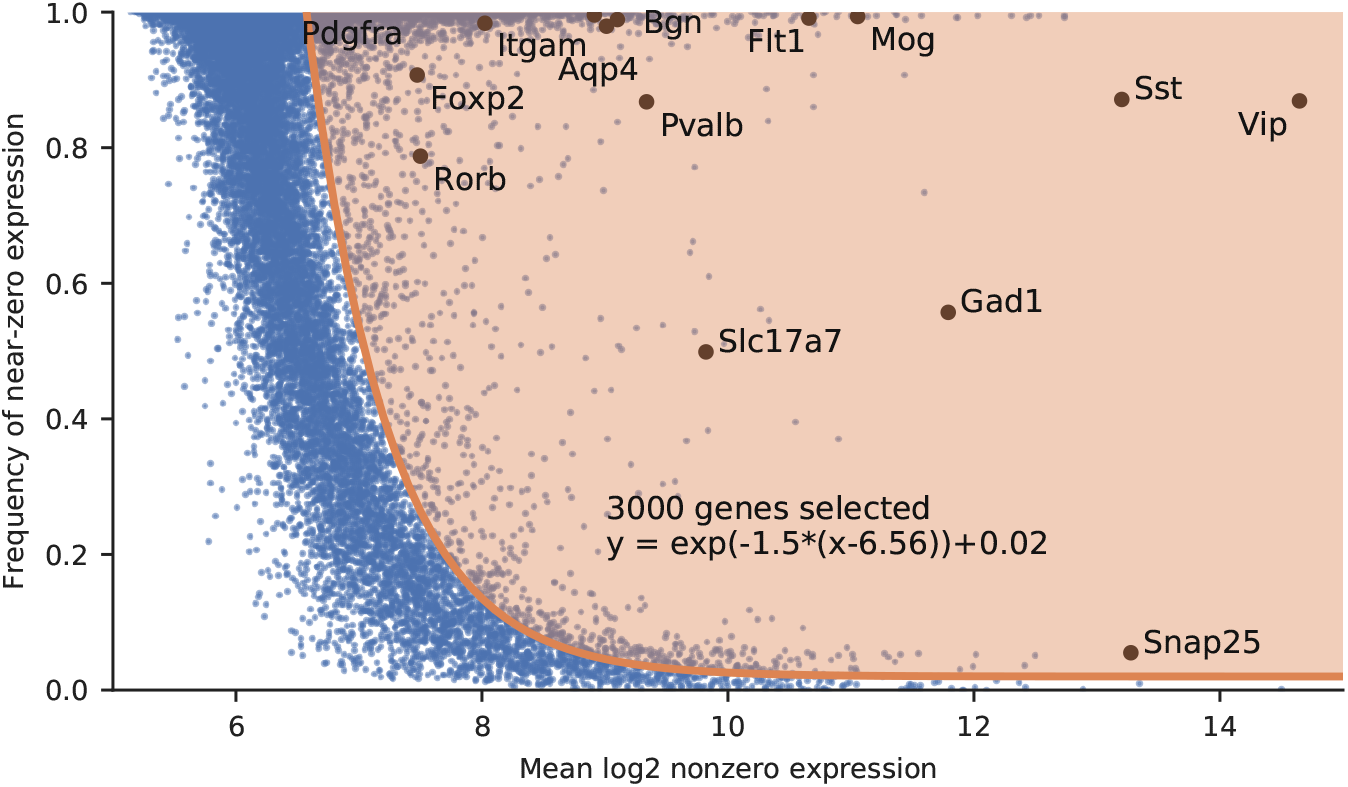
Feature selection. Our feature selection procedure illustrated for the Tasic et al. (2018) data set. Black dots show well-known marker genes, taken from Figure 1c of Tasic et al. (2016). Any good feature selection procedure should confidently select all of them.

